# Frizzled BRET sensors based on bioorthogonal labeling of unnatural amino acids reveal WNT-induced dynamics of the cysteine-rich domain

**DOI:** 10.1101/2021.05.01.442235

**Authors:** Maria Kowalski-Jahn, Hannes Schihada, Ainoleena Turku, Thomas Huber, Thomas P. Sakmar, Gunnar Schulte

## Abstract

Frizzleds (FZD_1-10_) comprise a class of G protein-coupled receptors containing an extracellular cysteine-rich domain (CRD) that binds lipoglycoproteins of the Wingless/Int-1 family (WNTs). Despite the prominent role of the WNT/FZD system in health and disease, our understanding of how WNT binding to the FZD CRD is translated into receptor activation and transmembrane signaling remains limited. Current hypotheses dispute the roles for conformational dynamics and the involvement of the linker domain connecting the CRD with the seven-helical transmembrane core of FZD. To clarify the mechanism of WNT binding to FZD and to elucidate how WNT/FZD complexes achieve signaling pathway specificity, we devised conformational FZD-CRD biosensors based on bioluminescence-resonance-energy-transfer (BRET). Using FZD engineered with N-terminal nanoluciferase and fluorescently-labeled unnatural amino acids in the linker domain and extracellular loop 3, we show that WNT-3A and WNT-5A induce similar CRD conformational rearrangements despite promoting distinct downstream signaling pathways, and that CRD dynamics are not required for WNT/β-catenin signaling. Thus, the novel FZD-CRD biosensors we report provide insights into the stepwise binding, activation and signaling processes in FZDs. The sensor design is broadly applicable to explore fundamental events in signal transduction mediated by other membrane receptors.

## Introduction

The class Frizzled of G protein-coupled receptors (GPCRs), also known as class F, comprises ten Frizzled subtypes (FZD_1-10_) and Smoothened (SMO)^1^. These cell surface membrane proteins share the structural hallmarks of GPCRs (extracellular N-terminus, seven membrane-spanning helices, TM1-7, connected *via* three extracellular and three intracellular loops, ECL1-3 and ICL1-3, and an intracellular C-terminus). FZDs also display a class-typical cysteine-rich domain (CRD) at an extended N-terminus, which is crucial for the engagement of their endogenous ligands with the receptor^2^.

While SMO regulates Hedgehog signaling, FZDs bind extracellular, secreted lipoglycoproteins of the Wingless/Int-1 family (WNTs) and mediate the vital effects of these 19 mammalian WNT paralogues on cellular proliferation, migration, differentiation, tissue polarity, tissue homeostasis and cancer development^3^. On the one hand, FZDs mediate signaling through dishevelled (DVL1-3)-dependent pathways resulting in either the stabilization and nuclear translocation of β-catenin or planar cell polarity-like signaling (PCP)^4–6^. On the other hand, they mediate DVL-independent signaling, through heterotrimeric G proteins^7–9^. Despite the enormous physiological relevance of the WNT/FZD signaling system in human health and disease, little is known about ligand/receptor selectivity, the molecular details that underlie receptor activation or the initiation of intracellular signaling. Thus, it remains unclear how distinct WNT/FZD complexes achieve pathway selectivity^10–12^.

Given the lack of structural information on WNT/FZD complexes and the fact that WNTs can bind to purified FZD-CRDs without the presence of the transmembrane core of the receptor^2^, uncertainty remains about the structural basis for agonist-induced, FZD-dependent signaling. As of today, two main models emerge, explaining how FZDs translate WNT/CRD association into different cellular signaling branches. One hypothesis is based on WNT-mediated crosslinking of FZDs with co-receptors (e.g., with low-density lipoprotein receptor-related protein 5 or 6 (LRP5/6), reversion-inducing cysteine-rich protein with Kazal motifs (RECK), tyrosine-protein kinase transmembrane receptor ROR2 or the adhesion GPCR ADGRA2 - also known as GPR124^13,14^), determining which intracellular transducer protein is recruited to subsequently convey the stimulus to the cell interior. This “signalosome” concept is widely accepted for signaling of FZDs through β-catenin, which depends on WNT-induced interaction of FZDs with LRP5/6^2,15–17^, and would allow for conformational flexibility of the extracellular linker region between the CRD and the receptor’s transmembrane core^18^. On the other hand, WNT-induced conformational changes in FZDs could also independently from WNT co-receptors promote distinct cellular signaling pathways^9,19–23^. The latter model mirrors the concept of how class A and B GPCRs activate, for instance, heterotrimeric G proteins in response to a ligand-induced opening of the intracellular receptor surface and subsequent GPCR/G protein coupling^24^. However, this process would require a constrained conformational mobility of the linker region to allow for mutual allosteric regulation of WNT binding and transducer coupling according to the ternary complex model^24,25^.

Although WNT-3A/β-catenin signaling occurs independently from heterotrimeric G proteins in human embryonic kidney cells (HEK293)^7^, it remains obscure whether and how the “signalosome” and “ternary complex” model are mechanistically intertwined in FZDs, hampering a rational development of FZD subtype-specific and intracellular pathway-selective drugs to treat WNT/FZD-dependent disorders such as colon and pancreatic cancer.

Recent studies provided intriguing but controversial insights into different aspects of the WNT/FZD system including the question of ligand/receptor selectivity and the underlying activation mechanism of FZDs^10,18,23,26^. For instance, while mutagenesis-based approaches suggested that FZD_5_ does not undergo conformational changes whilst signaling through DVL/β-catenin^18^, molecular dynamics (MD) simulations of FZD_4_^22^, as well as experiments with conformational FZD biosensors revealed structural flexibility and its importance for downstream signaling mediated by these receptors^9,19,21,23^. These conflicting findings highlight the need for advanced biophysical approaches and molecular tools to investigate WNT/FZD interaction and the conformational landscape of FZDs to better understand the FZD mode of action.

The assessment of conformational dynamics of GPCRs in living cells was facilitated by the design of fluorescence-(FRET) and bioluminescence-resonance-energy-transfer (BRET)-based biosensors already in the early 2000s^27,28^. Their development has not only allowed to monitor the structural rearrangements of various GPCRs – including FZD_5_ and FZD_6_^19,21^ – in real time and single cells, but refinements of the sensor design have further enabled the establishment of screening-compatible assay formats^29,30^. More recently, a distinct approach based on conformationally sensitive, circularly permutated fluorescent proteins (cpFPs), which were introduced to various GPCRs to monitor receptor activation in living animals^31–35^,provided unprecedented insights into WNT-induced conformational dynamics of FZDs^23^. While these conformational GPCR sensors exclusively detect the receptors’ structural dynamics at the intracellular side, several FRET-based GPCR sensors have been devised to study the extracellular conformational rearrangements of mainly class C GPCRs labeled using SNAP- and CLIP-tag technology^36–38^. In addition, genetic code expansion and labeling of unnatural amino acids enabled the investigation of tethered agonist (‘Stachel’) exposure in class adhesion GPCRs (aGPCRs)^39^.

In class F GPCRs, MD simulations based on an inactive SMO crystal structure (including a resolved CRD) revealed moderate CRD flexibility in absence of a ligand, which is further restrained upon cholesterol binding to the CRD^40^. Similarly, MD studies on an active SMO structure show only minor CRD movement when bound to cholesterol on the CRD and the receptor core allowing the receptor to adapt an active TM7 conformation to promote intracellular signaling^41^. However, the CRD-binding ligands of SMO and FZDs are distinctively different (cholesterol *versus* WNTs, respectively) and the linker domain between the CRD and the 7TM core of the receptor is notably shorter in SMO than in FZDs^1^ indicating that the functional dynamics of the CRD might also differ. FZD structures including a fully resolved CRD are not available and techniques to investigate extracellular dynamics of GPCRs in intact cells are limited by the size of fluorescent tags. Inserting, for instance, a fluorescent protein in one of the receptor’s extracellular loops would most likely impair a proper folding of the receptor and thereby abolish its expression at the cell surface or could sterically interfere with ligand/receptor association.

To overcome these limitations of conventional fluorescent labeling techniques, we linked small fluorescent probes to modified receptor residues using a minimally invasive labeling procedure based on genetic code expansion and incorporation of unnatural amino acids (uaa) serving as anchors for a subsequent bioorthogonal coupling reaction (strain-promoted inverse electron-demand Diels-Alder cycloaddition – SPIEDAC)^42–46^. With the aim to monitor and understand the WNT-induced dynamics of the extracellular CRD and its role for signal initiation and in order to avoid interference with WNT-binding upon incorporation of bulky tags, we developed a set of conformational biosensors based on BRET between the small NanoLuciferase (Nluc) and a fluorescently-labeled uaa. The minimal size of the energy acceptor maintained functionality and ligand binding to the receptor and allowed assessment of WNT-induced dynamics of the FZD extracellular domains in living cells, providing intriguing insights into the mechanistic and kinetic details of receptor activation preceding WNT/FZD signaling.

## Results

### MD simulations of a FZD_6_ model reveal CRD mobility

In order to assess the putative range of motion of the CRD relative to the receptor core, we employed first *in silico* approaches. In absence of a self-evident template for the extended linker region of FZDs, we prepared an inactive full-length model of FZD_6_ using the iTasser server^47–49^. FZD_6_ was selected for the modelling as it has a shorter extended linker sequence than other representatives of the four FZD homology clusters (i.e., FZD_4_, FZD_5_, and FZD_7_). The best-ranked model differs from the inactive SMO (PDB ID: 5L7D) only by this extended linker region (**Fig. 1a**) and was then used to initiate all-atom MD simulations (250 ns in seven independent replicas). In these simulations, the CRD probes a range of movements culminating to six main CRD orientations, which represent 62 % of the simulation trajectory (**Fig. 1c**). Of these, clusters 2 and 5 mark the conformational extremes of the CRD movements of the whole trajectory. The 7TM core including the disulfide-bond-stabilized part of the linker as well as the extended TM6 are stable throughout the simulation. Furthermore, the CRD remains stably folded and only its location relative to the receptor core is changing (**Fig. 1c, d**). Altogether, the MD data suggest that the CRD of FZD_6_ is capable to occupy distinct – conformationally restricted – orientations. To investigate whether these findings relate to the WNT binding, we set out to study how the overall orientation of the CRD relative to the receptor core is affected by WNT stimulation employing BRET technology.

**Figure 1.**
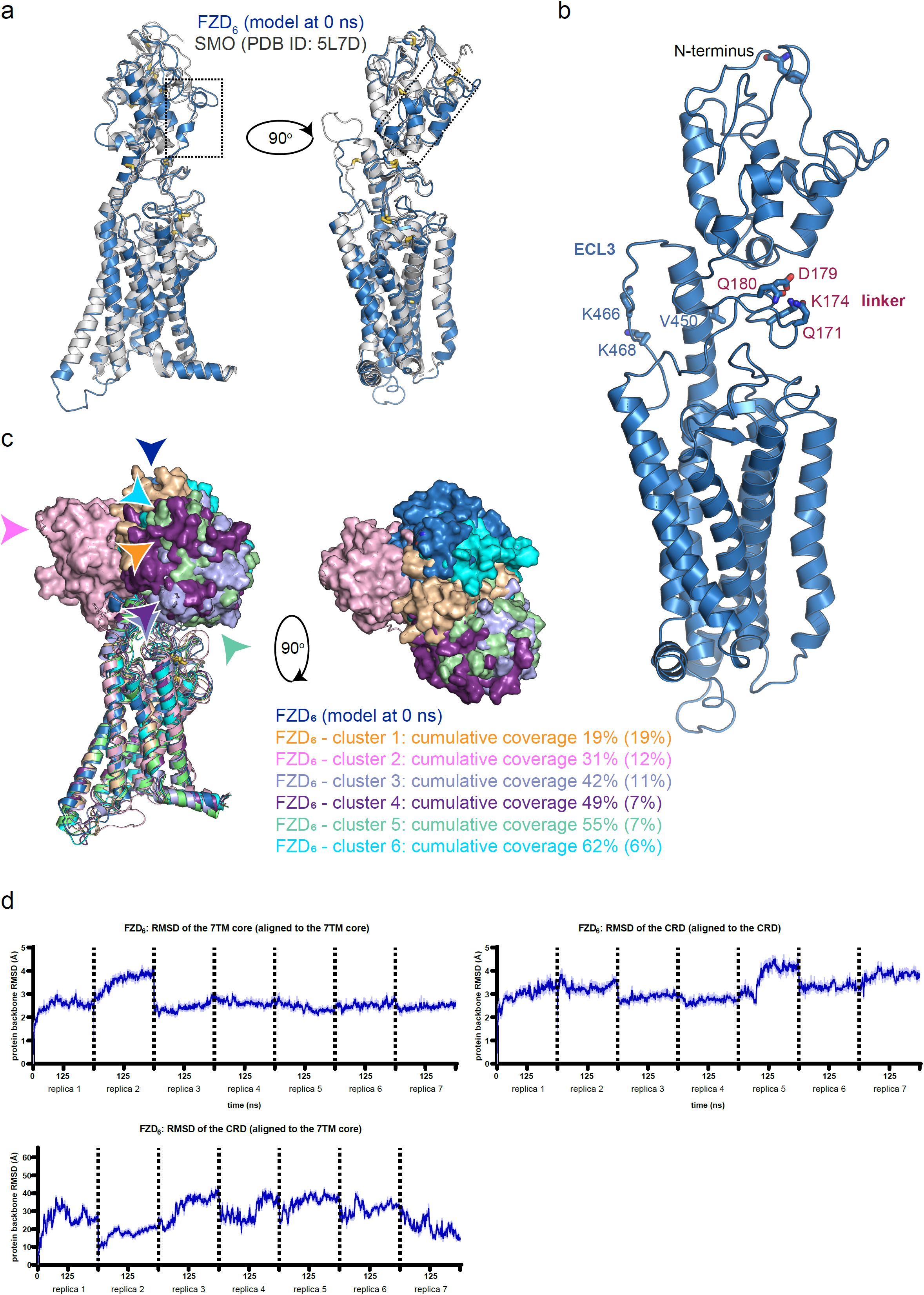
Molecular dynamics of FZD_6_ and receptor model with amber codon placement. a, Superimposition of the FZD_6_ model (blue) and inactive SMO (grey; PDB ID: 5L7D). Disulfide bridges are shown as sticks. The dotted rectangles mark the extended linker region, which differs between these two structures. b, A closer view to the FZD_6_ model. Amino acid residues selected for the point mutations are marked as sticks. Color code is as follows: red, oxygen; dark blue, nitrogen; light blue, carbon; yellow, sulfur. c, The CRD movements in the MD simulation. The receptor cores are shown as cartoon and CRDs as solid surfaces. Colored arrows mark the locations of the N-termini of each conformation cluster. Cumulative percentage of the time of coverage for each position cluster in the MD (absolute percentage for each cluster in parentheses). d, Protein backbone RMSDs of the receptor core (amino acids 160-511), and the CRD (amino acids 1-130) plotted as a continuous simulation trajectory. Dotted lines mark the independent simulation replicas. Thick blue traces indicate the moving average smoothed over a 2 ns window and thin traces the raw data. *CRD*, cysteine-rich domain, *ECL3*, extracellular loop 3, *TM*, transmembrane domain.

### Incorporation of TCO*K into class F GPCRs and bioorthogonal labeling

As representatives of class Frizzled, we chose FZD_6_ and FZD_5_, which belong to different homology clusters of class F^1^. The receptors were fused to an N-terminal Nluc epitope, following the 5-HT_3_ receptor signal peptide, and a C-terminal 1D4 epitope. By using the amber codon suppression technology, the unnatural amino acid (uaa) trans-cyclooct-2-ene-L-lysine (TCO*K) was introduced at distinct positions in the linker domain and the extracellular loop 3 (ECL3) of FZDs. We selected suitable positions for incorporation of TCO*K from the FZD_6_ simulation trajectory (**Fig. 1b**; **Suppl. Fig. 1**). The residues that were intended to be mutated were found to be in a distance towards the FZD’s N-terminus that allows BRET analysis.

For amber codon suppression, HEK293T cells were transfected with the amber-mutated receptor and the corresponding orthogonal suppressor tRNA/aminoacyl tRNA synthetase pair in presence of the uaa TCO*K. To boost the amber suppression, we introduced the respective amber mutant-bearing FZD into an expression plasmid carrying four repeats of the orthogonal suppressor tRNA (plasmid from Simon Elsässer, Addgene number: 140008). In addition, we defined a transfection ratio for the tRNA/FZD expression plasmid and the corresponding suppressor tRNA/aminoacyl tRNA synthetase plasmid^50^ (Addgene number: 140023) of 9:1 which results in an efficient uaa incorporation (**Fig. 2a**). Western blot analysis using mAb 1D4, which recognized a fused C-terminal epitope tag, showed that the amber codon suppression with the uaa TCO*K was efficient (**Suppl. Fig. 2a**). Cell surface expression of all TCO*K-incorporated FZD mutants was determined using whole-cell ELISA detecting the N-terminal Nluc epitope (**Suppl. Fig. 2b**). All mutants were expressed at the cell surface of HEK293T cells, averaging 31 % to 81 % compared with the respective wild-type receptor (**Suppl. Fig. 2b**).

**Figure 2.**
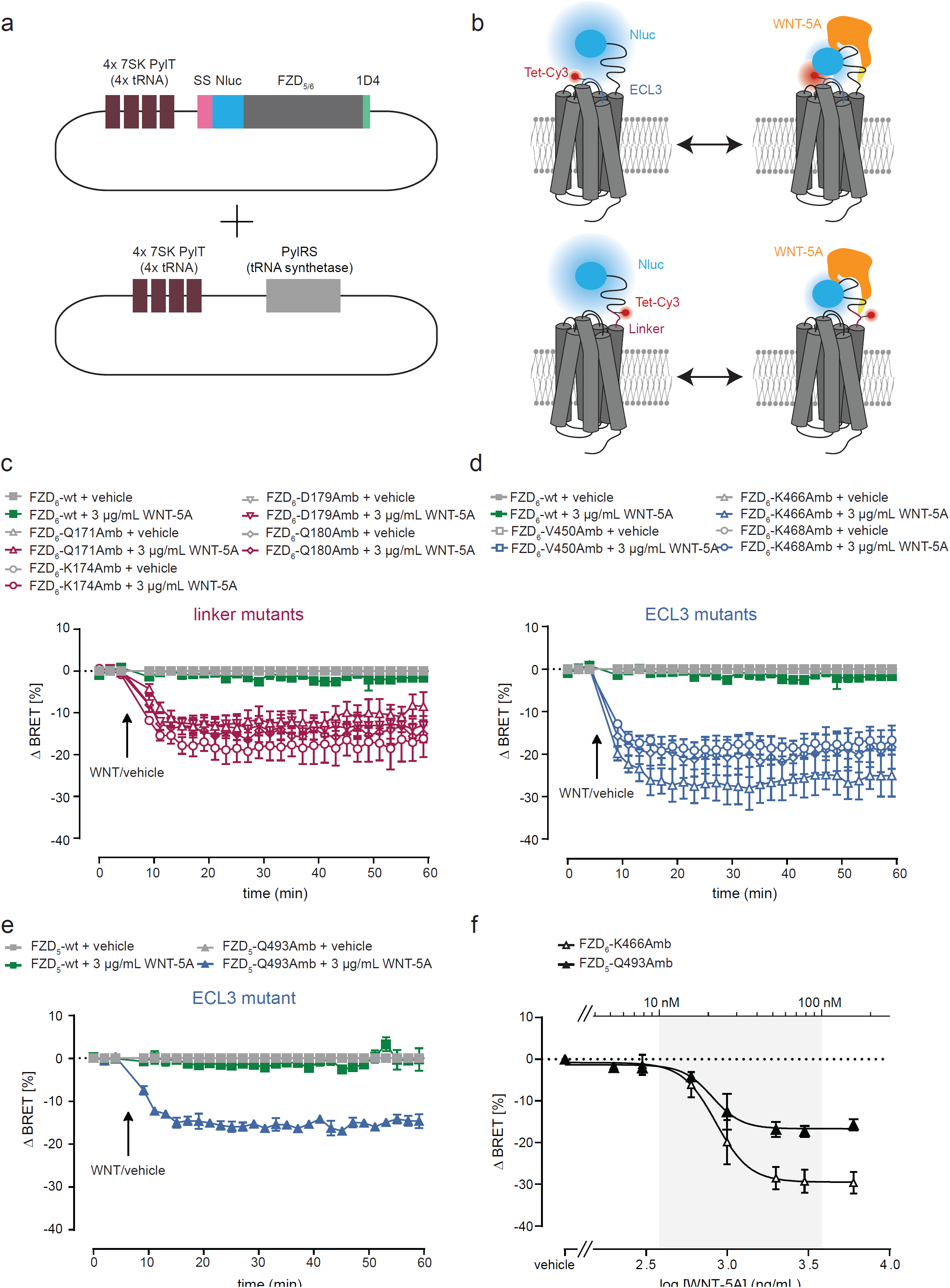
CRD movements in FZD_5_ and FZD_6_ in response to WNT-5A. a, Schematic depiction of cotransfected plasmids, one plasmid carrying four orthogonal tRNA repeats (4x 7SK PylT, Simon Elsässer, Addgene number: 140008) and amber-mutated FZD_5_ or FZD_6_ carrying an 5-HT_3_ receptor signal peptide, an N-terminal Nluc and a C-terminal 1D4 epitope tag, and the other plasmid carrying four orthogonal tRNA repeats and the tRNA synthetase^50^ (Addgene number: 140023). b, Schematic depiction of the CRD sensor design with the N-terminally Nluc-tagged FZD. Fluorescence labeling of residues in the linker (pink) or ECL3 (blue) region of the receptor with Tet-Cy3. c-d, BRET responses of FZD_6_ linker (c) and ECL3 (d) amber mutant CRD sensors upon 3 µg/mL WNT-5A treatment or vehicle control. The arrow indicates the time point of WNT or vehicle application. e, BRET responses of FZD_5_ ECL3 amber mutant CRD sensors upon 3 µg/mL WNT-5A treatment or vehicle control. The arrow indicates the time point of WNT/vehicle addition. f, Concentration-response curves of WNT-5A on HEK293T cells expressing FZD_5_-Q493Amb or FZD_6_-K466Amb. Concentrations-response curves are normalized to vehicle control. All experiments were performed in HEK293T cells cotransfected with the indicated FZD_5_ or FZD_6_ sensors and the orthogonal tRNA/synthetase pair. *SS*, signal sequence, *Nluc*, NanoLuciferase, *Amb*, amber mutant, *Tet*, tetrazine, *ECL3*, extracellular loop 3.

In a next step, the receptor mutants were expressed in HEK293T cells and living cells were labeled with the tetrazine (Tet)-bearing, membrane-impermeable fluorescent dye Tet-Cy3 (**Suppl. Fig. 3a**) using the SPIEDAC reaction. As an extension, also the membrane-permeable dye Tet-BODIPY-FL (BDP-FL) was used to label selected receptor mutants (**Suppl. Fig. 4a**). For quantification of the labeling efficiency of the different receptor mutants, a plate reader assay was used to detect fluorescence intensities (**Suppl. Fig. 3b and 4b**). While Tet-Cy3 specifically labeled HEK293T cells expressing receptor amber mutants exclusively in presence of the uaa TCO*K, Tet-BDP-FL showed more unspecific labeling properties (**Suppl. Fig. 4b**).

In general, cell surface expression levels of the receptor mutants correlated with the labeling efficiency.

### WNT-5A induced BRET changes in the FZD_5_ and FZD_6_ CRD sensors

We took advantage of the N-terminally fused Nluc and the fluorescent dye, incorporated site-specifically in the linker region or ECL3 of the receptor, to establish BRET biosensors that can detect WNT-induced conformational rearrangements of the CRD (**Fig. 2b**). Initially, we tested all FZD_6_ CRD sensors, comprising four receptor mutants in the linker region and three receptor mutants in the ECL3, in a ligand-free condition in terms of basal energy transfer between Nluc and the incorporated Cy3 or BDP-FL, respectively (**Suppl. Fig. 5**). Bioluminescence emission spectra of Nluc-tagged FZD_5_ and FZD_6_ CRD sensors, labeled with Tet-BDP-FL (green) or Tet-Cy3 (red), were recorded and normalized to the maximal Nluc emission. To exclude an unspecific fluorescent labeling of HEK293T cells expressing the receptor mutant sensors the bioluminescence signal obtained in cells expressing the wild-type receptor lacking incorporated TCO*K was subtracted. We obtained emission peaks at ∼510 nm for Tet-BDP-FL and ∼570 nm for Tet-Cy3, respectively, with highest peaks for the FZD_6_-K466Amb mutant.

After confirming the occurrence of intramolecular basal BRET for all mutants, we applied 3 µg/mL WNT-5A, which corresponds to 70.86 nM (molar mass of WNT-5A was taken from https://www.uniprot.org: 42,339 Da) to HEK293T cells expressing the different FZD CRD sensors and recorded the resulting BRET response of the Cy3-labeled FZD sensors over time. We detected dynamic WNT-induced BRET changes for all FZD_6_ linker (**Fig. 2c**) and ECL3 (**Fig. 2d**) mutant sensors with fitted maximal ΔBRET amplitudes varying between -11.61 ± 0.5116 % for the linker mutant FZD_6_-Q171Amb and -26.40 ± 0.7411 % for the ECL3 mutant FZD_6_-K466Amb (**Suppl. Table 1**).

We extended our studies to FZD_5_, which in contrast to FZD_6_ mediates both G protein- and WNT/β-catenin-dependent signaling^21,51^, and generated the ECL3 mutant FZD_5_-Q493Amb corresponding to FZD_6_-K466Amb. This FZD_5_ CRD sensors also displayed a WNT-induced conformational change of the CRD with a maximal ΔBRET amplitude of about -15.60 ± 0,2274 % (ΔBRET amplitudes in Fig. 2c-e were fitted with a plateau after decay equation, see **Suppl. Table 1**). Most importantly, the dynamic BRET signal illustrating the conformational change of the CRD towards the receptor core can be induced by WNT-5A in both the FZD_6_-K466Amb and the FZD_5_-Q493Amb sensor with similar EC_50_ values (850.3 ± 84.37 ng/mL for FZD_6_-K466Amb and 807.1 ± 94.57 ng/mL FZD_5_-Q493Amb; **Fig. 2f**). We tested also a labeling of the two ECL3 mutant sensors FZD_6_-K466Amb and FZD_5_-Q493Amb with Tet-BDP-FL and detected WNT-5A-induced BRET response for both sensors (**Suppl. Fig. 6a-c**). In contrast to Tet-Cy3–labeled sensors, Tet-BDP-FL-labeled sensors show positive BRET amplitudes, which can be explained by changes in relative dipole orientation as observed with other intramolecular GPCR biosensors^52–55^.

### WNT-3A-induced equivalent BRET changes in FZD_5_ and FZD_6_ CRD sensors

While WNT-5A is considered to initiate β-catenin-independent signaling, WNT-3A signals through β-catenin-dependent signaling including phosphorylation of low-density lipoprotein receptor-related protein 5 or 6 (LRP5/6), β-catenin stabilization and finally activating the TCF/LEF-dependent gene expression^10,56^ following the principles of the signalosome hypothesis.

We aimed to test whether 3 µg/mL WNT-3A, which corresponds to 76 nM (molar mass of WNT-3A was taken from https://www.uniprot.org/: 39,365 Da) is able to induce conformational changes not only in FZD_5_ but also in FZD_6_ known to mediate β-catenin-independent signaling (**Fig. 3a**). With both sensors, FZD_6_-K466Amb (**Fig. 3b**) and FZD_5_-Q493Amb (**Fig. 3c**), we were able to detect a WNT-3A-induced BRET response. Equal to WNT-5A, concentration-response curves of WNT-3A applied for 30 min revealed similar EC_50_ values for both receptor sensors (832.8 ± 24.83 ng/mL for FZD_6_-K466Amb and 869.6 ± 33.99 ng/mL for FZD_5_-Q493Amb; **Fig. 3d**). In fact, the maximal BRET responses of both the FZD_5_-Q493Amb and the FZD_6_-K466Amb sensor, for WNT-3A and WNT-5A are comparable to each other (**Suppl. Table 1**) and suggest that WNT-3A could activate FZD_6_ independently of LRP5/6 or more precisely, induces a conformational change of the CRD upon binding.

**Figure 3.**
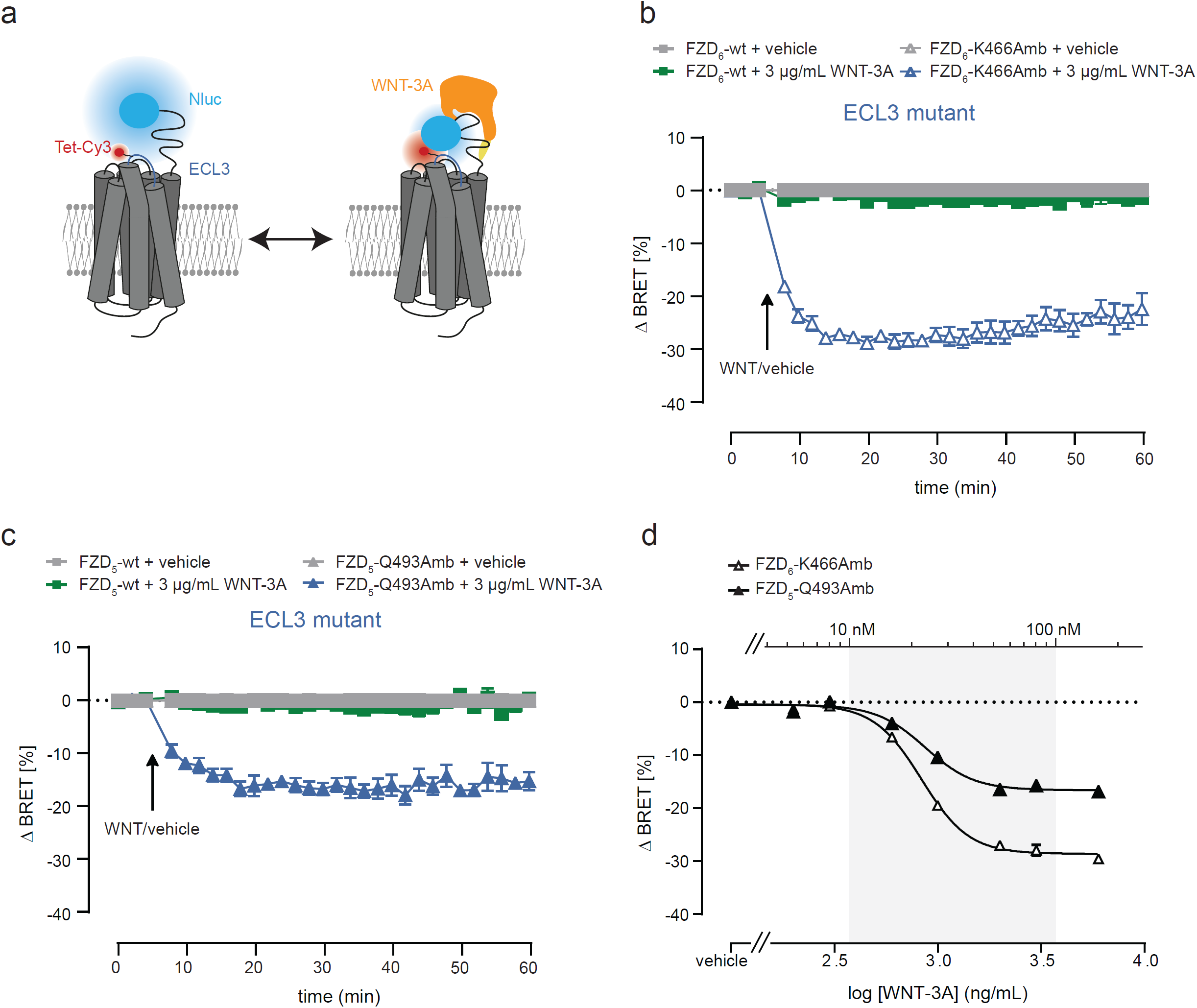
CRD movements in FZD_5_ and FZD_6_ in response to WNT-3A. a, Schematic depiction of the CRD sensor design with the N-terminally Nluc-tagged FZD. Fluorescence labeling of residues in the ECL3 (blue) region of the receptor with Tet-Cy3. b-c, BRET responses of FZD_6_ (b) or FZD_5_ (c) ECL3 amber mutant CRD sensors upon 3 µg/mL WNT-3A treatment or vehicle control. The arrow indicates the time point of WNT/vehicle addition. d, Concentration-response curves of WNT-3A on HEK293T cells expressing FZD_5_-Q493Amb or FZD_6_-K466Amb. Concentrations-response curves are normalized to vehicle control. All experiments were performed in HEK293T cells cotransfected with the indicated FZD_5_ or FZD_6_ sensors and the orthogonal tRNA/synthetase pair. *Nluc*, NanoLuciferase, *Amb*, amber mutant, *Tet*, tetrazine, *ECL3*, extracellular loop 3.

### Intramolecular versus intermolecular BRET responses

Little is known about FZD dimerization, but there is evidence that FZD dimerization through a CRD-CRD interaction contributes to WNT-induced β-catenin signaling as it was shown in Xenopus xFZD_3_^57^ or FZD_5_ and FZD_7_^58^. FZD_6_ exists as homodimer under basal conditions and undergoes dissociation and re-association upon WNT stimulation^59^.

The existence of FZD dimers could result in intermolecular “crosstalk” between Nluc and the FZD_5_ and FZD_6_ sensors of different monomers. In order to quantify the contribution of intermolecular BRET to the total BRET response of the FZD CRD sensors, we cotransfected a Nluc-tagged FZD wild type (wt), which is not *per se* able to act as a sensor, and a Nluc-lacking FZD amber mutant. While cotransfection of Nluc-FZD_5_-wt and the FZD_5_-Q493Amb did not result in a detectable WNT-induced BRET response compared to the BRET response detected with the intramolecular Nluc-FZD_5_-Q493Amb sensor (**Suppl. Fig. 7**), cotransfecting Nluc-FZD_6_-wt and the Nluc-lacking FZD_6_-K466Amb sensor did. The WNT-5A-induced BRET response obtained with the intermolecular BRET pair set-up, however, was smaller compared to the intramolecular Nluc-FZD_6_-K466Amb sensor (**Suppl. Fig. 8a, b**). The BRET response in the intermolecular sensor set-up indicated that a minor part of the total BRET change originates from agonist-induced FZD_6_ dimer dissociation. While these findings support the previous data on WNT-induced dimer dissociation^59^, the small contribution of the intermolecular BRET does not affect the conclusions about the intramolecular BRET changes with regard to CRD rearrangements. In order to provide further support of this assumption, we calculated the reaction constant k by fitting the BRET amplitudes for each of the experimental paradigms with intra- and intermolecular sensors. The higher k values for the intramolecular BRET changes compared to the intermolecular sensor argue for a chronological order of the events with faster extracellular conformational changes followed by FZD_6_ dimer dissociation (**Suppl. Fig. 8c**).

In addition, we made use of the FZD_6_ dimerization deficient triple Ala mutant D365A/R368A/Y369A^59^ by introducing the three Ala mutations into the Nluc-FZD_6_-K466Amb sensor, resulting in the Nluc-FZD_6_-K466Amb dimer mutant. The FZD_6_-K466Amb dimer mutant maintained cell surface localization (**Suppl. Fig. 9a**) and incubation with Tet-Cy3 showed a distinct fluorescence labeling capability (**Suppl. Fig. 9b**). The WNT-5A-induced conformational change resulted in BRET responses showing amplitudes of about half of the FZD_6_-K466Amb mutant (fitted plateau values: FZD_6_-K466Amb dimer mutant -10.22 ± 0.3197 % and FZD_6_-K466Amb -19.74 ± 0.6795 %) arguing for intramolecular BRET within one monomer between the Nluc fused to FZD’s N-terminus and the introduced fluorescent dye in the ECL3 (**Suppl. Fig. 9c**).

Thus, these dimer control experiments indicate that the BRET amplitudes as a consequence of WNT-induced extracellular conformational changes in FZD_6_ present a composite response of both intra- and intermolecular BRET events. In contrast, intermolecular BRET events have no impact on the FZD_5_ CRD sensor, which solely detects intramolecular BRET as a consequence of WNT-induced extracellular conformational changes.

### Kinetic insights into WNT-induced conformational changes of FZDs

WNT-induced FZD dynamics were previously investigated using intracellular fluorescence-based conformational sensors^23^. We aimed to compare the speed of the WNT-induced conformational rearrangements in distinct domains of FZDs by quantifying and comparing the reaction rates of the extracellular and intracellular conformational FZD sensors (**Fig. 4a**). Therefore, we stimulated HEK293T cells expressing the FZD_5_-Q493Amb CRD sensor (**Fig. 4b, c**) or the FZD_5_-cpGFP intracellular sensor (**Fig. 4b, d**) with 3 µg/µL WNT-3A and WNT-5A, and recorded the resulting BRET or fluorescence response of the two different sensors over time. The resulting reaction constant k was found to be significantly higher for the FZD_5_-Q493Amb CRD sensor for both, WNT-3A and WNT-5A (**Fig. 4b**). The higher reaction constant implies that the conformational changes at the extracellular part of FZD occur faster than the intracellular detected rearrangements, highlighting a sequence of events, where WNT-induced conformational changes in the extracellular domain precede those in the core of the receptor.

**Figure 4.**
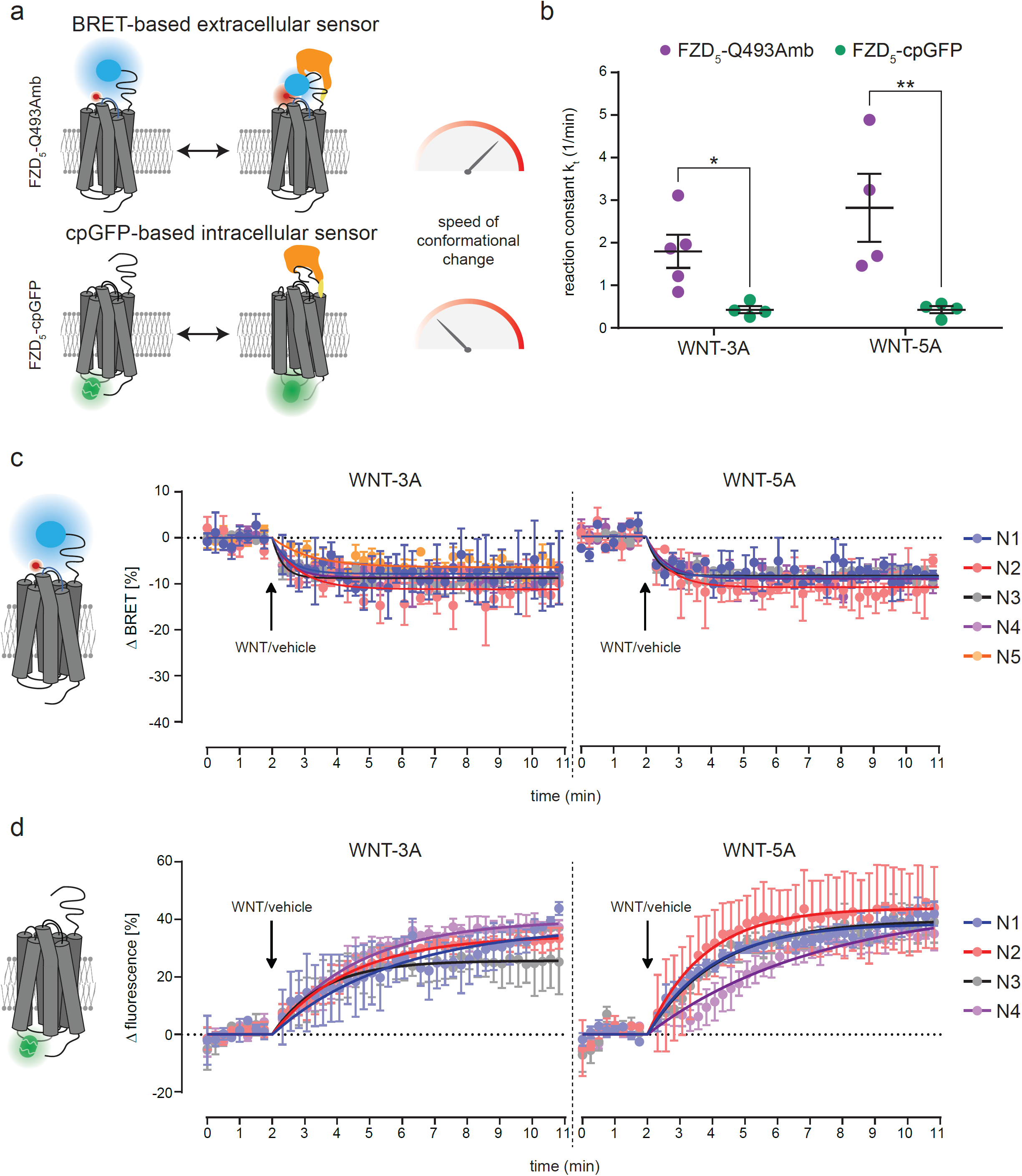
Kinetic insights into WNT-induced conformational changes in FZD_5_. a, Schematic depiction of the CRD sensor design for the BRET-based extracellular sensor FZD_5_-Q493Amb and the cpGFP-based intracellular sensor FZD_5_-cpGFP upon WNT binding. b, Reaction constant k of WNT-3A and WNT-5A induced BRET responses (FZD_5_-Q493Amb extracellular sensor) or fluorescence response (FZD_5_-cpGFP intracellular sensor) determined from fitted data in (c) and (d) using the plateau followed by one phase decay equation. Differences in WNT-3A- or WNT-5A-induced effects on FZD_5_-Q493Amb and FZD_5_-cpGFP were analyzed with two-way ANOVA followed by Fisher’s LSD post-hoc test. Significance levels are given as * (p < 0.05), and ** (p < 0.01). c, Kinetic fits of WNT-3A and WNT-5A induced BRET responses of HEK293T cells expressing FZD_5_-Q493Amb of five independent experiments (N1 - N5). d, Kinetic fits of WNT-3A and WNT-5A induced fluorescence responses of HEK293T cells expressing FZD_5_-cpGFP of four independent experiments (N1 – N4). *Amb*, amber mutant, *cpGFP*, circularly permutated green fluorescent protein.

### The role of LRP5/6 for WNT-induced conformational changes in the CRD

WNT-3A-mediated β-catenin signaling depends on a WNT-induced crosslinkage of FZDs with LRP5/6 (**Fig. 5a**). The crosslinkage can be blocked by the glycoprotein dickkopf-1 (DKK1), known to inhibit β-catenin-signaling by binding to the ectodomains of LRP5/6 and thereby preventing WNT engagement with LRP5/6 and subsequent FZD/LRP complex formation in signalosomes^60,61^.

**Figure 5.**
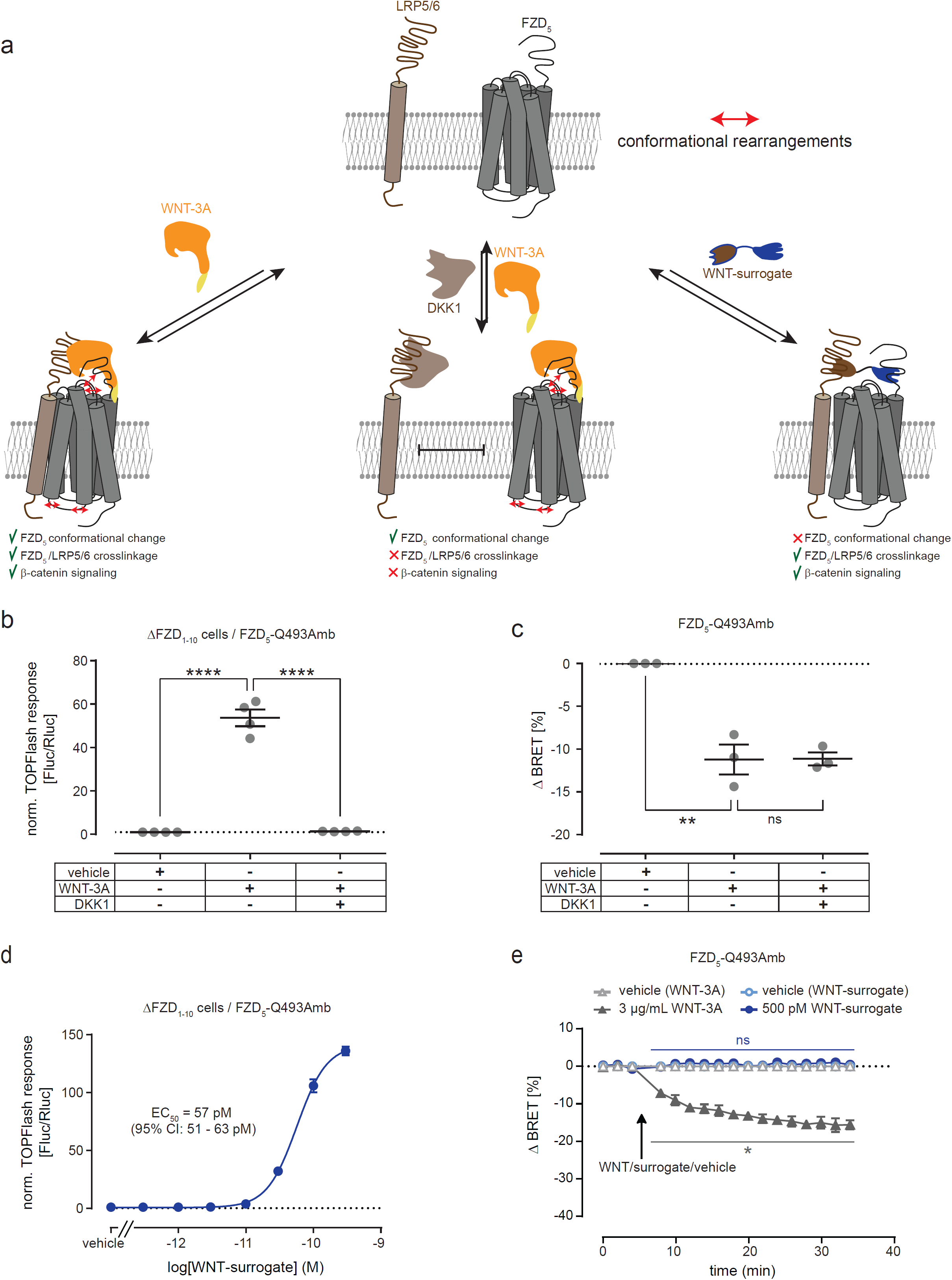
DKK1 and a WNT-surrogate to analyze the role of LRP5/6 for WNT-induced extracellular conformational changes. a, Schematic depiction of the initiation of WNT-induced β-catenin signaling. b, TOPFlash reporter gene response induced by 1 μg/mL WNT-3A in absence or presence of 1 μg/mL DKK1 in ΔFZD_1-10_ HEK293 cells transiently transfected with FZD_5_-Q493Amb. Data show mean ± s.e.m. of four independent experiments. c, Maximal BRET response induced by 3 μg/mL WNT-3A in absence or presence of 3 μg/mL DKK1 in ΔFZD_1-10_ HEK293 cells transiently transfected with FZD_5_-Q493Amb. Results in (b) and (c) were analyzed with one-way ANOVA followed by Tukey post-hoc test. Significance levels are given as ** (p < 0.01), **** (p < 0.0001), and *ns* (not significant). d, TOPFlash reporter gene response induced by increasing concentration of WNT-surrogate in ΔFZD_1-10_ HEK293 cells transiently transfected with FZD_5_-Q493Amb. The concentration-response curve of WNT-surrogate is normalized to vehicle control. e, BRET responses of the FZD_5_– Q493Amb sensor upon 3 µg/mL WNT-3A, 500 pM WNT-surrogate treatment or vehicle control four individual experiments. The arrow indicates the time point of WNT/WNT-surrogate or vehicle application. Differences between WNT-surrogate vehicle control and WNT-surrogate or WNT-3A vehicle control and WNT-3A-induced BRET responses were analyzed with multiple t-test followed by Holm-Sidak multiple comparison. Significance levels are given as * (p < 0.05), and *ns* (not significant). All experiments were performed in ΔFZD_1-10_ HEK293 cells or HEK293T cells cotransfected with FZD_5_-Q493Amb and the orthogonal tRNA/synthetase pair. *Amb*, amber mutant.

To strengthen the hypothesis that WNTs can promote conformational rearrangements in FZDs independently of co-receptors such as LRP5/6, we preincubated the FZD_5_-Q493Amb sensor with recombinant DKK1. While DKK1 inhibits WNT-3A induced FZD_5_/LRP5/6 signaling along the WNT/β-catenin axis (**Fig. 5b**), DKK1 preincubation did not affect WNT-induced conformational changes of the FZD CRD assessed by robust BRET responses (**Fig. 5c**) indicating that the WNT-induced conformational change of the CRD did not require FZD/LRP5/6-crosslinking.

In contrast, a surrogate WNT (named as WNT-surrogate), an artificial construct composed of a FZD- and a LRP5/6-binding moiety initiates β-catenin signaling through a forced crosslinkage of FZD with LRP5/6^62^. The WNT-surrogate applied to the FZD_5_-Q493Amb sensor yielded a concentration response curve for the induction of a TCF transcriptional response (TOPFlash assay) with an EC_50_ value of 57 pM (95 % CI: 51 – 63 pM; **Fig. 5d**). However, even at a ten times higher concentration (500 pM) the WNT-surrogate was not able to induce any BRET responses arguing for the absence of extracellular conformational changes that can be detected by the FZD_5_ CRD sensor (**Fig. 5e**). These findings indicate that WNT-3A induces FZD_5_/LRP5/6 crosslinkage and extracellular conformational changes in FZDs, but that the latter are not required to initiate β-catenin signaling.

## Discussion

The WNT/FZD system represents a primary component of myriad vital biological processes in human physiology and disease. Binding of WNTs to the CRD of FZDs constitutes the initial step of WNT morphogen signaling^2^, triggering diverse intracellular signaling cascades through crosslinking of FZDs with WNT co-receptors (most prominently with LRP5/6)^2,15–18^ and by inducing intramolecular, conformational dynamics in FZDs independently from co-receptor interaction^23^. Seminal work in models of *Drosophila melanogaster* has allowed to infer WNT/FZD interaction from intracellular signaling-dependent readouts^26,63–65^ and more recent studies with conformational receptor biosensors have provided valuable insights into WNT-induced structural dynamics at the intracellular parts of FZDs^19,21,23^. However, how WNT engagement with the extracellular CRD is mechanistically connected to these intracellular receptor conformational changes or FZD/WNT-co-receptor crosslinking remained unclear. The flexibility of the CRD relative to the core of FZDs and its relevance for signal initiation has been a matter of debate and it remains unclear how the information flow from WNT binding to the CRD is transduced to the core of the receptor provided that the linker domain is freely flexible.

Our study provides structural and kinetic insights into this core event of WNT/FZD signaling. Here, we describe the development and validation of optical biosensors that unveil extracellular conformational rearrangements in FZDs in real-time in living cells. These extracellular conformational biosensors rely on BRET between N-terminally fused Nluc and a fluorescent dye, incorporated in the receptor’s linker region or ECL3 using a minimally invasive technique based on site-directed insertion of uaas and bioorthogonal coupling chemistry in live cells minimizing interference with receptor functionality and ligand/receptor interaction. One important mechanistic finding of our study is that the FZD CRD rearrangements detected by our BRET sensors are not required to initiate β-catenin signaling. Our experiments with a WNT-surrogate known to mediate β-catenin signaling through crosslinkage of FZDs with LRP5/6^66^ showed that (i) WNT-surrogate/FZD_5_/LRP5/6 assembly does not provoke conformational changes in the FZD_5_ CRD sensor and (ii) β-catenin-dependent signaling can be mediated by the FZD_5_ biosensor without extracellular conformational changes. These observations support previous notions of a receptor tyrosine kinase (RTK)-like functionality of FZDs that exclusively relies on clustering FZD and LRP5/6 in a signalosome^18,23^.

Furthermore, we discovered that WNT-3A and WNT-5A induced very similar BRET response at saturating concentrations, arguing for analogous CRD rearrangement upon ligand binding. This finding is somewhat surprising in light of the distinct modes of action and intracellular signaling pathways initiated by these two endogenous FZD ligands (FZD/co-receptor crosslinking and β-catenin-dependent signaling by WNT-3A *versus* FZD conformational changes and β-catenin-independent signaling by WNT-5A). This analogy poses the question of whether WNT-3A, concurrent to co-receptor-dependent, RTK-like signal propagation, mediates similar functionalities of FZD_5/6_ as WNT-5A by inducing conformational changes in the receptor. Supporting this hypothesis, WNT-3A mediates phosphorylation of extracellular signal-regulated kinases 1 and 2 (ERK1/2) in primary mouse microglia^56^ and regulates GTPase activity in platelets^67^, processes that are often associated with GPCR/G protein signaling. Interestingly, WNT-3A signaling through the WNT/β-catenin pathway and β-catenin-independent signaling to ERK1/2 occurs simultaneously in mouse primary microglia, albeit with slightly similar kinetics, regulating the proinflammatory status of these brain macrophages. Our current data underline that the same WNT can elicit both signalosome-dependent and FZD conformation-dependent signal initiation providing a molecular and mechanistic basis for FZD functional selectivity.

In addition, our study sheds light on the stepwise processes occurring between WNT-CRD binding and FZD/transducer coupling. Employing a set of distinct BRET- and fluorescence-based assays, we show that CRD movements take place before the rearrangement of the receptors’ intracellular domains is initiated, and, in the case of FZD_6_, before ligand-induced dissociation of receptor homodimers. These observations highlight the extracellular CRD movement in FZDs as a prime event underlying WNT/FZD signaling and confirm – using an unprecedented live cell biosensor system – the central modulatory role of the CRD in class F GPCRs^40^. It remains to be defined, what molecular movement in fact determines the agonist-induced changes in BRET using the FZD CRD sensors. In an attempt to better understand the consequences of ligand association with the CRD, we overlaid the WNT structure with the different CRD position clusters extracted from the MD simulations. This schematic overlay (**Suppl. Fig. 1d**) suggests that one mechanism resulting in a ΔBRET could be the restriction of the range of motion of the CRD through WNT binding, which in addition could be accompanied by ligand-induced conformational rearrangements in the receptor core. More experiments are required to dissect these structural details of FZD activation.

In summary, the sensor design described here presents a universal approach to develop cell-based optical probes for membrane-spanning proteins, including other classes of GPCRs, receptor-tyrosine kinases or cytokine receptors, aiding in the mechanistic exploration of fundamental biological processes underlying ligand initiation, kinetics of receptor conformational changes and cellular signaling in general.

## Materials and Methods

### Computational studies

The FZD_6_ model (amino acids 22-511) was built using iTasser server^47–49^. The protocol selected ten templates for the modelling: 2ZIY (squid rhodopsin), 4JKV (inactive ΔCRD-SMO), 4ZWJ (rhodopsin-arrestin complex), 5L7D (inactive SMO), 5NDD (protease-activated receptor-2), 6BD4 (ΔCRD-FZD_4_), 6FJ3 (parathyroid hormone 1 receptor), 6LW5 (formyl peptide receptor 2), 6ME6 (melatonin receptor MT2), and 6WW2 (ΔCRD-FZD_5_). After visual examination of the produced five models, two models – model 1 and model 3 which both were close to our previous inactive ΔCRD-FZD_6_ model^23^ (RMSD 2.4 and 3.2 Å, respectively) and had similarly folded CRDs than FZD_5_ (PDB ID: 5URZ) and FZD_7_ (PDB ID: 5T44) – were picked for a preliminary MD study (ca. 150 ns of production simulation after the equilibration protocol, for details see below). Only model 1 remained stable in the preliminary simulation, and was thus selected for the further MD study.

The MD simulations were run using GROMACS 2020.3^68^. FZD_6_ was oriented by aligning it to the FZD_4_ from OPM database^69^ and embedded in the POPC lipid bilayer (150 lipids / leaflet) by CHARMM-GUI server^70^ with TIP3p water molecules and 0.15 M NaCl. The system was minimized for approximately 2,000 steps and then equilibrated with gradually decreasing position restraints on protein and lipid components. In the last 50 ns of the equilibration run, the harmonic force constants of 50 kJ mol^-1^ nm^-2^ were applied on the protein atoms only.

Seven (250 ns) independent isobaric and isothermic (NPT) ensemble production simulations were initiated using the CHARMM36m force field^71^ and a 2 fs time step. First, replica 1 was run starting from the equilibrated structure and random velocities. Then, six other replicas were simulated starting from the snapshots of the replica 1 at time points t=0 ns, t=50 ns, t=100 ns, t=150 ns, t=200 ns, and t=250 ns and random velocities. The temperature at 303.15 K was maintained with Nose-Hoover thermostat^72^ and the pressure at 1 bar with Parrinello-Rahman barostat^73^. Potential-shift-Verlet was used for electrostatic and van der Waals interactions with 12 Å cut-off, and the bonds between hydrogen and other atoms were constrained by the LINCS algorithm^74^. The data were analyzed using VMD (visualization, measurement of RMSDs and distances, and RMSD clustering)^75^ and visualized in PyMol. For RMSD clustering, the size of the trajectory was reduced to contain one frame per every 10 ns of the simulation, the frames were superimposed on the 7TM core of the receptor at the first frame of the first replica, similarity cut-off was set to 5 Å and maximum number of clusters to 30. The cluster seeds were used as the representative models of each cluster. The MD data will be deposited to GPCRmd (an open-access MD database for GPCRs; www.gpcrmd.org).

### Cloning of FZD constructs

The synthetic gene constructs for FZD_5_ and FZD_6_ were designed based on the amino acid sequences of the full-length receptor lacking the native signal peptide (residues 27-585 of Uniprot Q13467 (FZD5_HUMAN) and residues 19-706 of Uniprot O60353 (FZD6_HUMAN)), and codon-optimized for expression in human cells using the GeneArt online tool while avoiding a set of motifs corresponding to several restriction sites (NheI, HindIII, NcoI, EcoRI, SbfI, MfeI, KpnI, NotI, and XbaI). We extended the 5’-end of the genes with the nucleotide sequence 5’-AAG CTT GCC GCC ACC ATG GCG CTG TGT ATC CCT CAA GTT CTG CTG GCC CTG TTC CTG AGC ATG CTG ACA GGA CCT GGC GAG GGC TAC CCT TAC GAT GTG CCT GAC TAC GCC GAA TTC GCT CCT GCA GGG AGT CAA TTG-3’ that adds the restriction site HindIII used for cloning and the protein sequence MALCIPQVLLALFLSMLTGPGEGYPYDVPDYAEFAPAGSQL to the N-terminus of the receptor. This sequence is derived from the mouse serotonin 5-HT_3_ receptor cleavable signal sequence carrying the R2A mutation to enable usage of a strong Kozak consensus sequence (GCCGCCACCATGG, start codon underlined) and an hemagglutinin (HA)-tag (YPYDVPDYA) followed by a linker EFAPAGSQL corresponding to a nucleotide sequence with EcoRI, SbfI, MfeI and MlyI sites^76^. We extended the 3’-end with the sequence 5’-GGT ACC GCC TCC TCG GAT GAG GCC AGC ACA ACC GTG TCT AAG ACC GAG ACA TCT CAG GTG GCC CCT GCC TAA GCG GCC GCT CTA GA-3’ containing a KpnI site at the 5’-end and NotI and XbaI sites at the 3’-end. The C-terminal 18 residue long sequence is the rhodopsin 1D4 mAb epitope tag^77^. The constructs were synthesized and cloned into a plasmid derived from pcDNA3.1(+) modified to eliminate an internal MfeI site.

All FZD_5_ and FZD_6_ amber mutants were generated using the GeneArt™ Site-directed Mutagenesis System (Thermo Fisher Scientific) with the following primers: FZD_6_-Q171Amb forward 5’-AAA ACA TCT GGC GGC TAG GGC TAC AAG TTC C-3’, FZD_6_-Q171Amb reverse 5’-GGA ACT TGT AGC CCT AGC CGC CAG ATG TTT T-3’, FZD_6_-K174Amb forward 5’-GGC GGC CAG GGC TAC TAG TTC CTG GGC ATC G-3’, FZD_6_-K174Amb reverse 5’-CGA TGC CCA GGA ACT AGT AGC CCT GGC CGC C-3’, FZD_6_-D179Amb forward 5’-AAG TTC CTG GGC ATC TAG CAG TGC GCC CCT CCA-3’, FZD_6_-D179Amb reverse 5’-TGG AGG GGC GCA CTG CTA GAT GCC CAG GAA CTT-3’, FZD_6_-Q180Amb forward 5’-TTC CTG GGC ATC GAT TAG TGC GCC CCT CCA T-3’, FZD_6_-Q180Amb reverse 5’-ATG GAG GGG CGC ACT AAT CGA TGC CCA GGA A-3’, FZD_6_-V450Amb forward 5’-TGG GAG ATC ACA TGG TAG TCC GAC CAC TGC AG-3’, FZD_6_-V450Amb reverse 5’-CTG CAG TGG TCG GAC TAC CAT GTG ATC TCC CA-3’, FZD_6_-K466Amb forward 5’-TGT CCA TAC CAG GCC TAG GCC AAA GCC AGA C-3’, FZD_6_-K466Amb reverse 5’-GTC TGG CTT TGG CCT AGG CCT GGT ATG GAC A-3’, FZD_6_-K468Amb forward 5’-TAC CAG GCC AAG GCC TAG GCC AGA CCT GAG CTG-3’, FZD_6_-K468Amb reverse 5’-CAG CTC AGG TCT GGC CTA GGC CTT GGC CTG GTA-3’, FZD_5_-Q493Amb forward 5’-GGC CAC GAT ACA GGC TAG CCT AGA GCC AAG C-3’ and FZD_5_-Q493Amb reverse 5’-GCT TGG CTC TAG GCT AGC CTG TAT CGT GGC C-3’.

The tRNA expression vector was a kind gift from Simon Elsässer (Addgene number: 140008). For cloning the BRET extracellular sensors, a NanoLuciferase (Nluc) tag in combination with FZD_5_ or FZD_6_ was subcloned into the tRNA expression vector using the NEBuilder® HiFi DNA Assembly kit (New England biolabs). Briefly, the tRNA expression vector was digested with NheI and BamHI. The Nluc tag obtained from the Nluc-FZD_6_ construct^19^ was cloned N-terminally of FZD_5_ or FZD_6_ with the following primers: Nluc forward 5’-TCC AAG CTG TGA CCG GCG CCT ACT CTA GAG CTA GCC ACC ATG CGG CTC TGC-3’, Nluc reverse 5’-TGC AGG AGC GAA TTC CGC CAG AAT GCG TTC GCA C-3’, FZD_5/6_ forward 5’-GAA CGC ATT CTG GCG GAA TTC GCT CCT GCA GGG AGT C-3’ and FZD_5/6_ reverse 5’-GCA GAC AGC GAA TTA ATT CCA GCG GCC GCG GAT CCG GCC GCT TAG GCA GGG GC-3’. For the dimer control construct, FZD_5_-Q493Amb or FZD_6_-K466Amb was subcloned without the N-terminally Nluc tag into the tRNA expression vector using the NEBuilder® HiFi DNA Assembly kit with the following primers: FZD_5/6_ forward 5’-GCT GTG ACC GGC GCC TAC TCT AGA GCT AGC GCC GCC ACC ATG GCG CTG-3’ and FZD_5/6_ reverse 5’-CAG CGA ATT AAT TCC AGC GGC CGC GGA TCC GGC CGC TTA GGC AGG GGC-3’. The tRNA expression vector was digested with NheI and BamHI. The triple alanine mutant D365A/R368A/Y369A mutant (called the FZD_6_-K466Amb dimer mutant) was cloned using the GeneArt™ Site-Directed Mutagenesis PLUS kit (Thermo Fisher Scientific) into the tRNA-Nluc-FZD6-K466Amb plasmid with the following primers: D365A/R368A/Y369A forward 5’-GCC TGT ACG ATC TGG CCG CCA GCG CGG CCT TTG TGC TCC TGC C-3’ and D365A/R368A/Y369A reverse 5’-GGC AGG AGC ACA AAG GCC GCG CTG GCG GCC AGA TCG TAC AGG C-3’.

All cloned constructs were verified by sequencing (eurofins Genomics).

### Cell culture, transfection and treatments

HEK293T cells cultured in Dulbecco’s modified Eagle’s medium with 1 % penicillin/streptomycin, and 10 % fetal bovine serum (all from Thermo Fisher Scientific) in a humidified 5 % CO_2_ incubator at 37 °C. Cells were transfected 24 h after seeding with Lipofectamine 2000 according to supplier’s information (Invitrogen). Absence of mycoplasma contamination was routinely confirmed by PCR using 5′-GGCGAATGGGTGAGTAACACG-3′ and 5′-CGGATAACGCTTGCGACTATG-3′ primers detecting 16 S ribosomal RNA of mycoplasma in the media after 2–3 days of cell exposure.

Treatments were done 48 h after transfection using the following agents: recombinant WNT-3A (R&D systems, 5036-WN-010), recombinant WNT-5A (R&D systems, 645-WN-010), surrogate WNT surrogate-Fc fusion protein (U-Protein Express B.V., N001^66^), dickkopf-1 (DKK1, R&D systems, 5439-DK-010/CF). The lyophilized preparations of recombinant WNTs and DKK1 were dissolved in 0.1 % BSA/DPBS and stored at 4 - 8 °C for maximum 4 weeks. Lyophilized WNT-surrogate was dissolved in 0.1 % BSA/DPBS and stored at 4 - 8 °C for maximum 4 weeks. Prior treatment in cell-based experiments, 10-fold compound dilutions were prepared in Sigmacote (Merck)-precoated transparent 96-well plates to avoid adsorption of recombinant proteins to plastic surfaces. All pipetting steps were performed with Sigmacote-precoated pipette tips.

### Immunoblotting

The day prior transfection, 100,000 HEK293T cells / well were seeded in 24-well plates. The cells were transfected with 0.45 μg of the indicated constructs and 0.05 µg of tRNA/synthetase^50^ (Addgene number: 140023; control conditions were balanced with pcDNA) per well and were cultured in the absence or presence of 0.1 mM trans-cyclooct-2-ene-L-lysine (TCO*K, Sichem, SC-8008). Because the FZD_5_-Q493Amb construct expressed weakly, 300,000 HEK293T cells / well were seeded in 12-well plates. The cells were transfected with 0.9 μg of the FZD_5_-Q493Amb construct or 0.18 µg FZD_5_-wt/ 0.72 µg pcDNA3.1 and 0.1 µg of tRNA/synthetase (control conditions were balanced with pcDNA). Cells were lysed 48 h after transfection in 2x Laemmli buffer containing 200 mM dithiothreitol (DTT, Merck). Lysates were sonicated and separated by SDS-PAGE/immunoblotting using 7,5 % gels. Transfer to a polyvinylidene difluoride membrane was done with the Trans-Blot® Turbo Transfer System (Bio-Rad). After transfer, membranes were incubated in 5 % low-fat milk/TBS-T (25 mm Tris- HCl, 150 mm NaCl, 0.05 % Tween 20, pH 7.6) and subsequently in primary antibodies overnight at 4 °C. The next day, the membranes were washed four times in TBS-T, incubated with goat anti-mouse or goat anti-rabbit secondary antibody conjugated to horseradish peroxidase (Thermo Fisher Scientific, 1:5,000), washed, and developed using Clarity™ Western ECL Substrate (Bio-Rad) according to the supplier’s information. Primary antibodies were as follows: anti-1D4 (National Cell Culture Center, mouse, 1:1,000), anti-Nluc (R&D systems, MAB100261-SP, mouse, 1:500) and GAPDH (Cell Signaling Technology, 2118, rabbit, 1:4,000).

### Whole-cell ELISA

For quantification of cell surface receptor expression, 15,000 HEK293T cells were plated in 96-well plates precoated with 0.1 mg/mL poly-D-lysine (PDL). Next day, cells were transfected with 0.09 μg of the indicated constructs and 0.01 µg of tRNA/synthetase and were cultured in the absence or presence of 0.1 mM TCO*K. After 48 h, cells were incubated with an anti-Nluc antibody (R&D systems, MAB100261-SP, mouse, 1:500) in 1 % BSA/DPBS for 1 h at 4 °C. Following incubation, cells were washed five times with 0.5 % BSA/DPBS and probed with a horseradish peroxidase–conjugated goat anti-mouse antibody at a 1:2,500 dilution in 1 % BSA/DPBS for 1 h at 4 °C. The cells were washed five times with 0.5 % BSA/DPBS, and 100 μl of the peroxidase substrate 3,3′,5,5′-tetramethylbenzidine (Merck) was added (30 min at room temperature). After acidification with 100 μl of 2 M HCl, the absorbance was read at 450 nm using a POLARstar Omega plate reader (BMG Labtech).

### Live cell imaging

The day prior transfection, 15,000 HEK293T cells / well were seeded into PDL-precoated black 96-well glass bottom plates. The cells were transfected with 0.09 μg of the indicated constructs and 0.01 µg of tRNA/synthetase and were cultured in the absence or presence of 0.1 mM TCO*K. 48 h after transfection, cells were washed in DPBS, kept for 2 h in DMEM to remove remaining TCO*K and were labeled with 1 µM Tet-Cy3 (Jena Bioscience, CLK-014-05) or Tet-BDP-FL (Jena Bioscience, CLK-036-05) for 30 min. Cells were washed again with DPBS and kept for additional 30 min in DMEM. Finally, DMEM was exchanged with 0.1 % BSA/HBSS and cells were imaged using a Zeiss LSM880 confocal microscope.

### Fluorescence labeling measurements

For quantification of the fluorescence labeling of the receptor mutants, 15,000 HEK293T cells were plated in black PDL-precoated 96-well plates. Next day, cells were transfected with 0.09 μg of the indicated constructs and 0.01 µg of tRNA/synthetase and were cultured in absence or presence of 0.1 mM TCO*K. 48 h after transfection, cells were washed in DPBS, kept for 2 h in DMEM and were labeled with 1µM Tet-Cy3 (Jena Bioscience, CLK-014-05) or Tet-BDP-FL (Jena Bioscience, CLK-036-05) for 30 min. Cells were washed again with DPBS and kept for additional 30 min in DMEM. Finally, DMEM was exchanged with 0.1 % BSA/HBSS and fluorescence intensities were read using a POLARstar Omega plate reader (BMG Labtech) equipped with filters for Cy3 (Ex. 544 nm / Em. 590 nm) and BDP-FL (Ex. 485 nm / Em. 520 nm).

### Nluc-FZD BRET measurements

For BRET measurements with the Nluc-FZD_5/6_ CRD sensors, 15,000 HEK293T cells were plated in PDL-precoated black 96-well plates. Next day, cells were transfected with 0.09 μg of the indicated constructs and 0.01 µg of tRNA/synthetase and were cultured in presence of 0.1 mM TCO*K. 48 h after transfection, cells were washed in DPBS, kept for 2 h in DMEM, and were labeled with 1 µM Tet-Cy3 or Tet-BDP-FL for 30 min. Cells were washed with DPBS and kept for additional 30 min in DMEM. Next, cells were again washed with DPBS and incubated with 90 μL of a 1/1000 dilution of furimazine stock solution (Promega) in 0.1 % BSA/HBSS. After 5 min of incubation, the basal BRET ratio was measured in three consecutive reads and 10 μL of a 3 μg/mL WNT-3A or WNT-5A solution (in 0.1 % BSA/HBSS) or vehicle control was applied per well. Subsequently, the BRET ratio was recorded for additional 25–60 min. For experiments with a higher temporal resolution, eight baseline BRET reads were recorded within 2 min prior manual addition of compounds or vehicle control, followed by at least 40 reads. All experiments were conducted using a CLARIOstar plate reader equipped with monochromators to separate Nluc (450/80 nm) and BDP-FL (520/40 nm) and Cy3 (580/30 nm), respectively.

### FZD-cpGFP fluorescence measurements

HEK293A cells stably expressing FZD_5_-cpGFP^23^ were seeded at a density of 80,000 cells/well onto PDL-pre-coated, black-wall, black-bottomed 96-well plates. 24 h later, all wells were washed with HBSS and incubated with 0.1 % BSA/HBSS. Baseline fluorescence was recorded in eight consecutive reads within 2 min, 10 μl of 10-fold WNT solution or vehicle control was applied per well and the resulting fluorescence intensity was recorded for additional 40 reads. All experiments were conducted using a CLARIOstar plate reader (BMG Labtech, Ortenberg, Germany) equipped with filters to excite cpGFP (470/15 nm) and record its emission intensity (515/20 nm). 40 flashes were applied per data point.

### TOPFlash reporter gene assay

ΔFZD_1-10_ HEK293 cells (400,000 cells / mL) were transfected in suspension with 400 ng M50 Super 8xTOPFlash, 100 ng pRL-TK Luc, 450 ng Nluc-FZD_5_-Q493Amb and 50 ng tRNA/synthetase per mL cell suspension, were supplemented with 0.1 mM TCO*K and seeded onto PDL-precoated white-wall, white-bottomed 96-well plates (50,000 cells / well). 24 h after transfection, cells were washed with 100 µl HBSS and incubated for 4 h in 72 µl / well of FBS-reduced (0.5 %) DMEM supplemented with 10 nM C59. Thereafter, 8 µl of 10 µg/mL recombinant WNT-3A and/or DKK1 (in 0.1 % BSA/HBSS), varying concentrations of WNT-surrogate or the respective vehicle controls were added. 24 h after stimulation, cells were washed with HBSS and lysed in 30 µl of Promega’s dual luciferase passive lysis buffer. Subsequently, 20 µl luciferase assay reagent (LARII) was added to each well and β-catenin-dependent firefly luciferase (Fluc) intensity was measured using a CLARIOstar microplate reader (580/80 nm; 1 sec integration time). Next, 20 µl Stop&Glo Reagent was added to quantify Renilla luciferase (Rluc) emission intensity (480/80 nm; 1 sec integration time) to control for variations in cell number and transfection efficiency.

### Statistical analysis

All immunoblot experiments are representative of three independent experiments. Statistical and graphical analysis was done using GraphPad Prism 9 software. For analyzing the surface expression, the basal absorbance detected in pcDNA-transfected HEK293 cells was subtracted from all data and mean values were normalized to wild-type FZD_5_ or FZD_6,_ which was set to 100 %. Differences among the TCO*K-untreated and -treated groups were analyzed by one-way ANOVA with uncorrected Fisher’s Least Significant Difference (LSD) test. Significance levels are given as: * (p < 0.05), ** (p < 0.0196), *** (p < 0.001), and **** (p < 0.0001). All data points represent normalized values, each performed in triplicates. Bars show means ± s.e.m. of three to four independent experiments.

Differences in fluorescence labeling among the TCO*K-untreated and -treated groups were analyzed by one-way ANOVA with uncorrected Fisher’s Least Significant Difference (LSD) test. Significance levels are given as: * (p < 0.05), ** (p < 0.01), *** (p < 0.001), and **** (p < 0.0001). All data points represent mean values, each performed in triplicates. Bars show means ± s.e.m. of three to six independent experiments.

BRET ratios were interpreted as acceptor emission/donor emission. At least three individual BRET reads were averaged before ligand/vehicle application. For quantification of ligand-induced changes, ΔBRET was calculated for each well as a percent over basal BRET. Subsequently, the average ΔBRET of the vehicle control was subtracted. All data points represent mean values, each performed in triplicates or quadruplicates, ± s.e.m. of three to five independent experiments. For analyzing the ΔBRET data, the WNT-induced averaged BRET responses were fitted with a plateau followed by one phase decay equation. Data from concentration-response experiments were fitted using a four-parameter fit. All data are represented as mean ± s.e.m. of at least three independent experiments. Data from TOPFlash experiments were expressed as ratios of Fluc over Rluc luminescence intensity to correct for distinct transfection efficiencies in the different samples. The resulting TOPFlash ratios were subsequently normalized for the average TOPFlash ratio of vehicle-treated wells to express WNT-induced changes as increases over baseline.

## Acknowledgments

Most importantly, we thank Simon Elsässer and Birthe Meineke for advice and tools to set-up the orthogonal labeling of FZDs. We also thank Anna Krook for access to the CLARIOstar plate reader and Benoit Vanhollebeke for the ΔFZD_1-10_ HEK293 cells. The work was supported by grants from Karolinska Institutet, the Swedish Research Council (2017-04676; 2019-01190), the Swedish Cancer Society (CAN2017/561, 20 0264P), the Novo Nordisk Foundation (NNF20OC0063168, NNF17OC0026940, NNF19OC0056122), Wenner-Gren Foundations (UPD2018-0064; UPD2019-0193), and the German Research Foundation (DFG; KO 5463/1-1, 427840891). Computational resources were provided by the Swedish National Infrastructure for Computing (SNIC) - National Supercomputer Centre (NSC) in Linköping and KTH Royal Institute of Technology (PDC) in Stockholm (SNIC 2020/5-500).

## Author contributions

MKJ and GS conceived and designed the study. MKJ, HS performed the wet lab experiments. AT performed the *in silico* work. TH and TPS provided know how and synthetic gene design. MKJ, HS, AT, GS designed and prepared the figures. MKJ, HS, AT, GS wrote the manuscript. TS commented and contributed to the manuscript writing. GS supervised and coordinated the project.

## Data and materials availability

Data supporting the findings of this manuscript are available from the corresponding author upon reasonable request. A reporting summary for this article is available as a Supplementary Information file. Expression vectors used and created for this work can be obtained through an MTA. MD simulations will be accessible on www.gpcrmd.org.

