## Supplementary Material for "Frizzled BRET sensors based on bioorthogonal labeling of unnatural amino acids reveal WNT-induced dynamics of the cysteine-rich domain"

[¤] Current affiliation: Orion Pharma R&D, P.O. Box 65 (Orionintie 1), FI-02101 Espoo, Finland

Supplementary Table 1. **Fitted ΔBRET amplitudes of all $FZD_5$ and $FZD_6$ sensors.**

| | ΔBRET amplitudes in % (plateau followed by one phase decay equation ± s.e.m.) | | |
|---|---|---|---|
| | **3 µg/mL WNT-5A** | | **3 µg/mL WNT-3A** |
| | Tet-Cy3 labeling | Tet-BDP-FL labeling | Tet-Cy3 labeling |
| **$FZD_6$** | | | |
| Wild type | -2.323 ± 1.022 | 0.1041 ± 0.04673 | -1.657 ± 0.2094 |
| Q171Amb | -11.61 ± 0.5116 | | |
| K174Amb | -17.91 ± 0.654 | | |
| D179Amb | -13.24 ± 0.4227 | | |
| Q180Amb | -14.88 ± 0.455 | | |
| V450Amb | -18.16 ± 0.4078 | | |
| K466Amb | -26.40 ± 0.7441[#] | 6.371 ± 0.1065 | -26.30 ± 0.3770[#] |
| K468Amb | -20.22 ± 0.8163 | | |
| **$FZD_5$** | | | |
| Wild type | -0.8425 ± 0.2687 | -1.103 ± 0.2306 | -0.8637 ± 0.2133 |
| Q493Amb | -15.60 ± 0.2274[##] | 3.70 ± 0.1005 | -16.01 ± 0.3105[##] |

[#], [##], no significant difference between effect of WNT-5A and WNT-3A on $FZD_6$-K466Amb and $FZD_5$-Q493Amb as analyzed with Student's unpaired t-test.

### Supplementary Figure 1

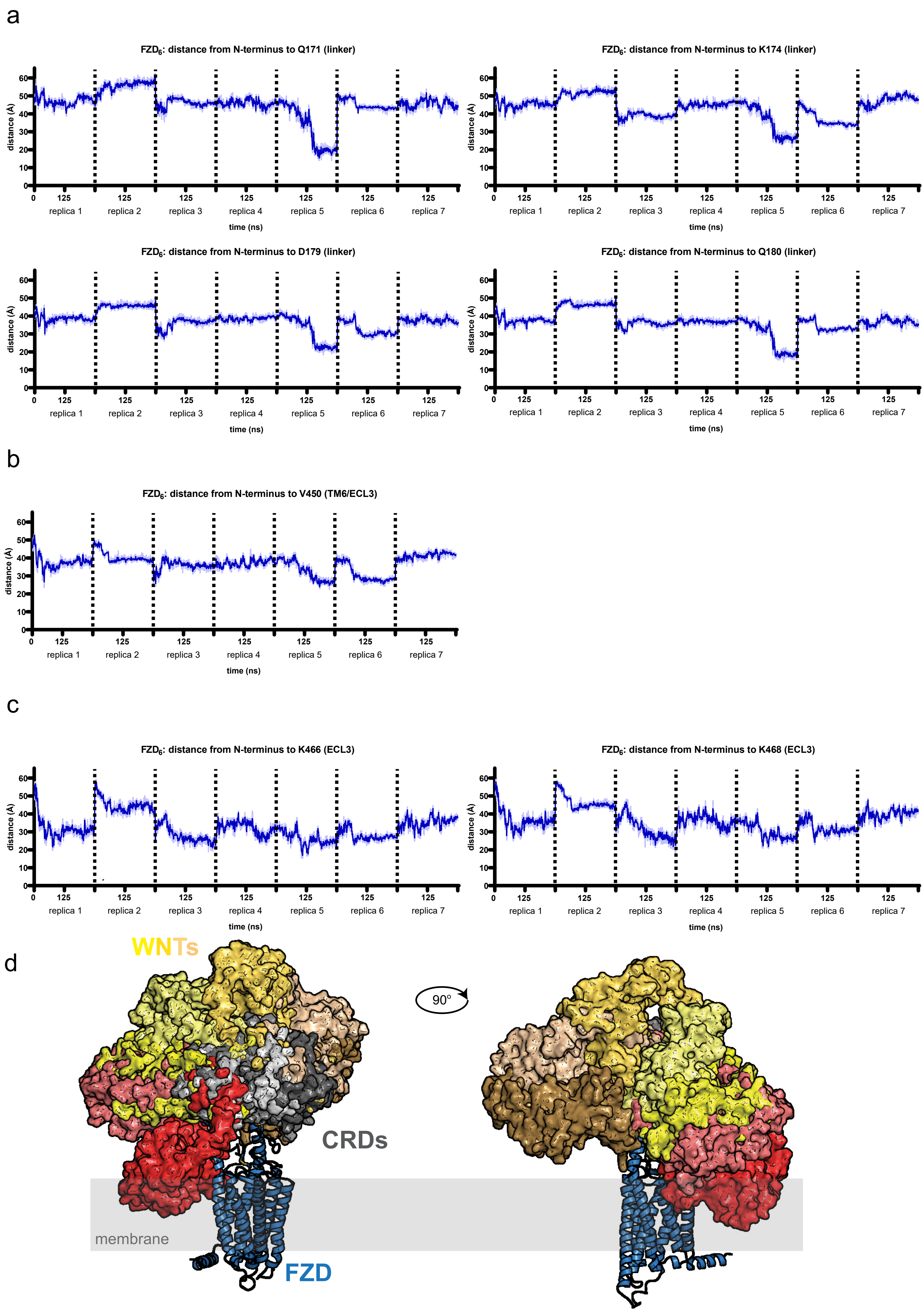

Supplementary Figure 1. **Distances between the unnatural amino acid (uaa) mutants and the N-terminus of the $FZD_6$ model throughout the simulation trajectory.** a, Mutants located at the disulfide-bridge-stabilized linker region. b, Mutants located at the extracellular extension of TM6. c, Mutants located at the ECL3. The distances are measured between the β-carbons of the residues picked for the point mutation and the N-terminal nitrogen atom and plotted as a continuous trajectory. Dotted lines mark the independent simulation replicas, thick blue traces indicate the moving average smoothed over a 2 ns window and thin traces the raw data. d, Schematic and hypothetical presentation of WNT binding to seven CRD clusters obtained from the MD simulations. The model of xWNT-8 from the xWNT-8-m$FZD_8$-CRD structure (PDB ID: 4F0A) was superimposed to CRDs of seven $FZD_6$ clusters. One $FZD_6$ core is coloured in blue, CRDs of the different $FZD_6$ cluster are coloured in different shades of grey, WNTs are coloured in shades of yellow or red depending on its position regarding relative to the cell membrane (= red means WNT clashes with cell membrane). Approximate localization of the plasma membrane is indicated with grey shading. *CRD*, cysteine-rich domain, *ECL3*, extracellular loop 3, *TM*, transmembrane domain.

### Supplementary Figure 2

a

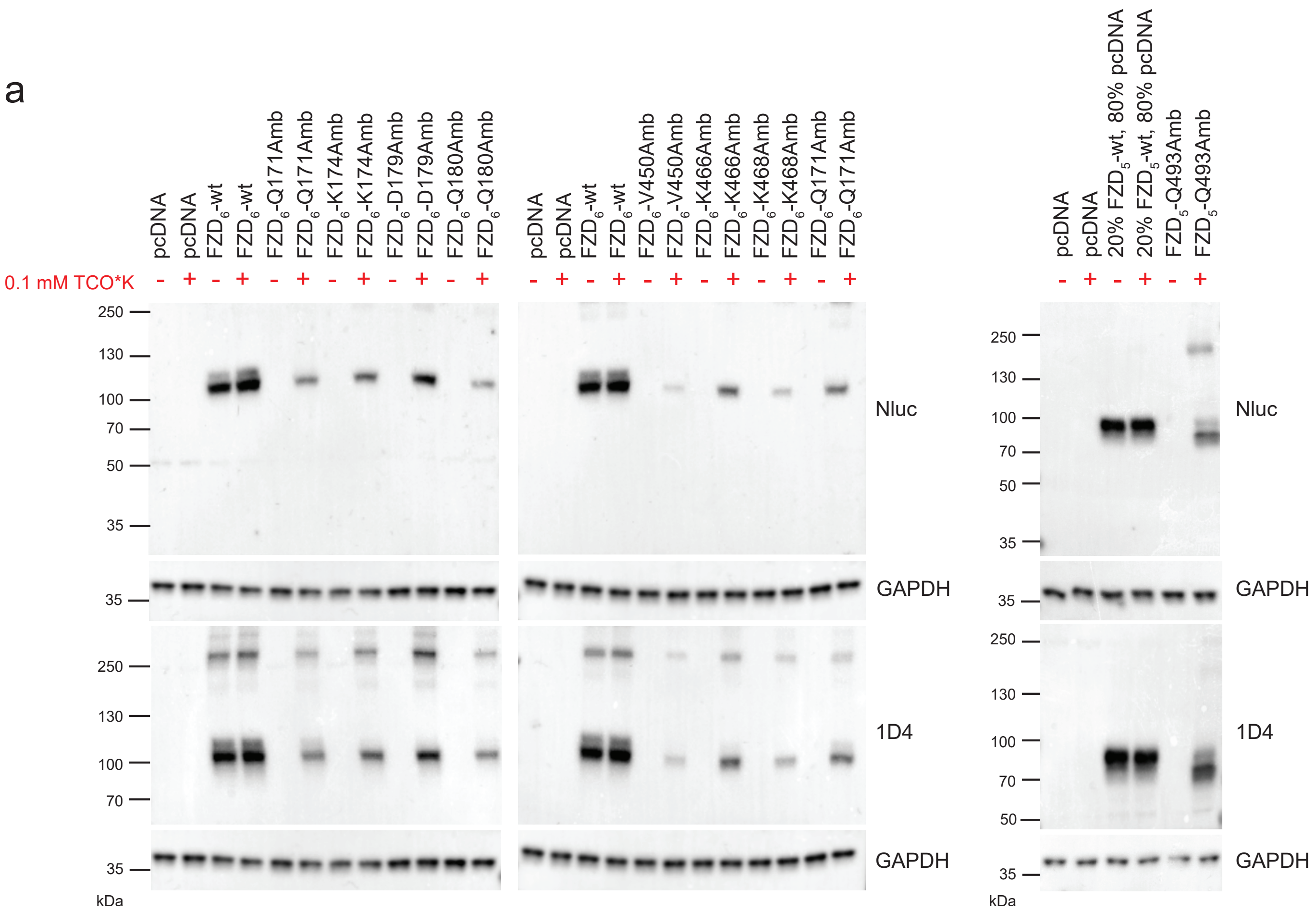


b

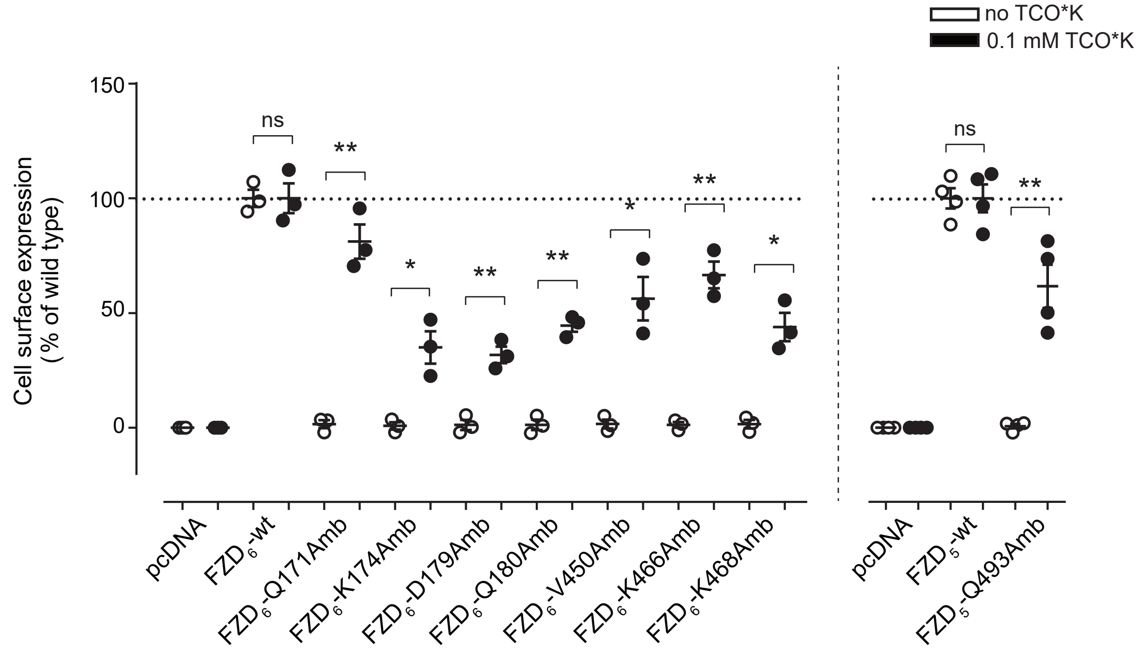

Supplementary Figure 2. **Characterization of $FZD_5$ and $FZD_6$ uaa mutants.** a, Amber suppression of HEK293T cells cotransfected with pcDNA3.1 (control), wt Nluc-$FZD_5$ or $FZD_6$ and the indicated amber mutants in absence (-) or presence (+) of 0.1 mM TCO*K. Cells were lysed and analyzed by immunoblotting using anti-1D4 (for detection of the full-length receptor) and anti-Nluc (detection of the N-terminal Nluc tag) antibodies. Anti-GAPDH served as a loading control. Note that the amount of $FZD_5$-wt was reduced to 20 % of the amount of transfected $FZD_5$-Q493Amb mutant ($FZD_5$-wt was balanced with pcDNA). b, Receptor surface expression of HEK293T cells transiently transfected with pcDNA3.1 or the indicated $FZD_5$ and $FZD_6$ constructs in absence (-) or presence (+) of 0.1 mM TCO*K was quantified by whole-cell ELISA using an antibody against the N-terminal Nluc tag. Data show mean ± s.e.m. of three to four individual experiments performed in triplicates. Background fluorescence detected in pcDNA-transfected HEK293T cells was subtracted from all data, and mean values were normalized to wt $FZD_5$ or $FZD_6$ surface expression. Results were analyzed with one-way ANOVA and uncorrected Fisher's LSD post-hoc test. Significance levels are given as * ($p < 0.05$), ** ($p < 0.01$), and *ns* (not significant). *Amb*, amber mutant, *TCO*K*, TCO-Lysine, *wt*, wild type, *uaa*, unnatural amino acid.

### Supplementary Figure 3

a

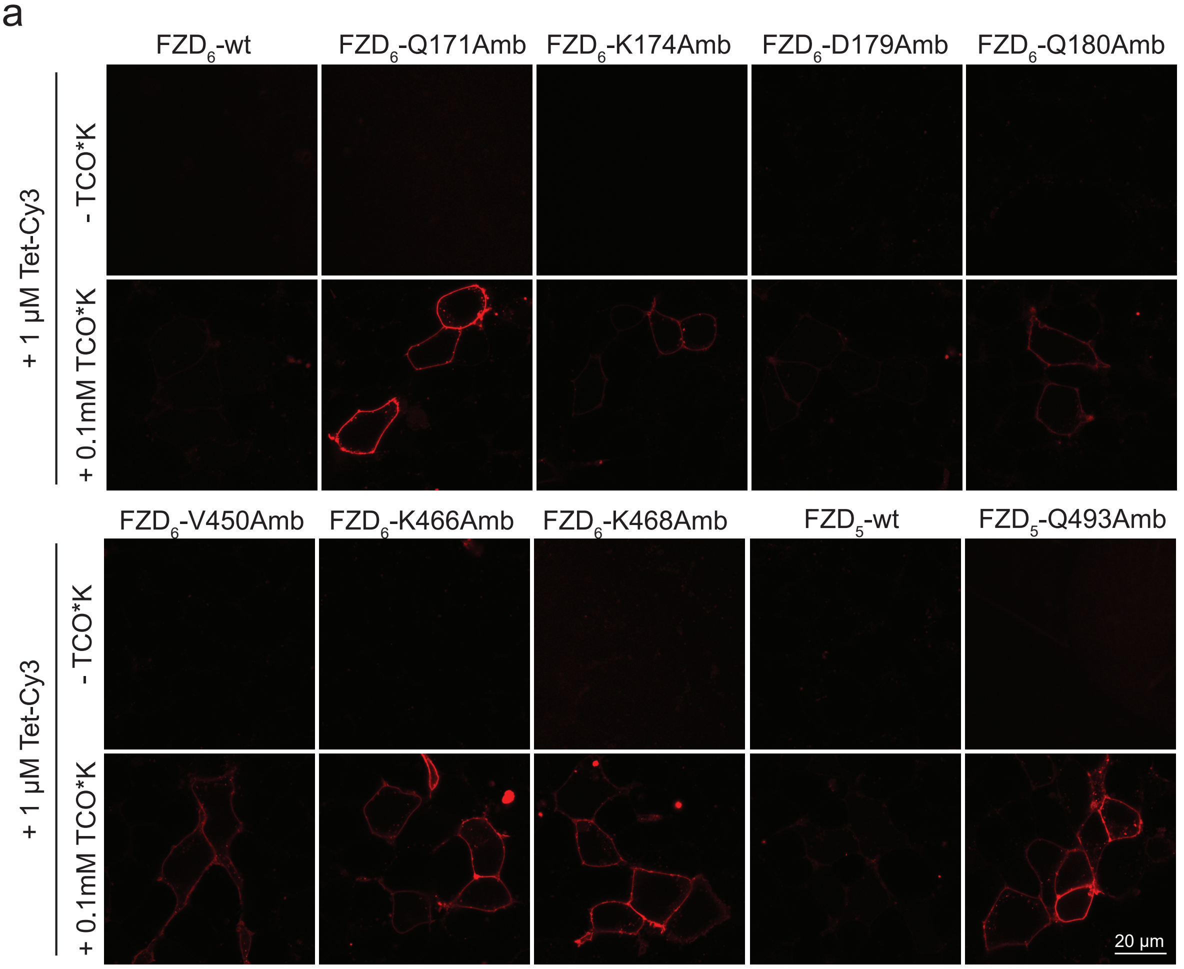


b

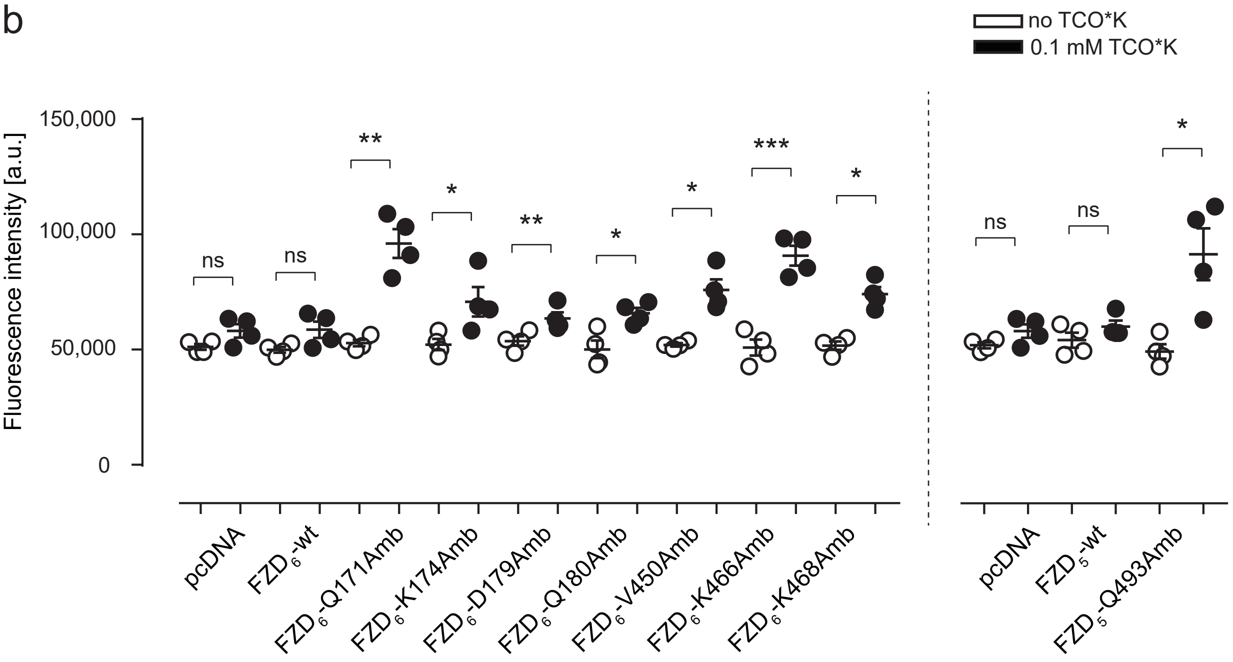

Supplementary Figure 3. **Fluorescence labeling of $FZD_5$ and $FZD_6$-TCO*K-incorporated mutants with Tet-Cy3.** a, Representative confocal images of HEK293T cells transiently cotransfected with $FZD_5$-wt, $FZD_6$-wt or the indicated amber mutants and the orthogonal tRNA/synthetase pair in absence (-) or presence (+) of 0.1 mM TCO*K. TCO*K-incorporated mutants were labeled with 1 μM of the cell membrane-impermeable fluorescent dye Tet-Cy3. *Scale bar*, 20 μm. b, Fluorescence intensities of HEK293T cells transiently cotransfected with $FZD_5$-wt, $FZD_6$-wt or the indicated amber mutants and the orthogonal tRNA/synthetase pair in absence (-) or presence (+) of 0.1 mM TCO*K after labeling with Tet-Cy3 measured in a plate reader assay. Data show mean ± s.e.m. of four individual experiments performed in triplicates. Results were analyzed with one-way ANOVA and uncorrected Fisher's LSD post-hoc test. Significance levels are given as * ($p < 0.05$), ** ($p < 0.01$), *** ($p < 0.001$), and *ns* (not significant). *Amb*, amber mutant, *TCO*K*, TCO-Lysine, *Tet*, Tetrazine, *wt*, wild type.

### Supplementary Figure 4

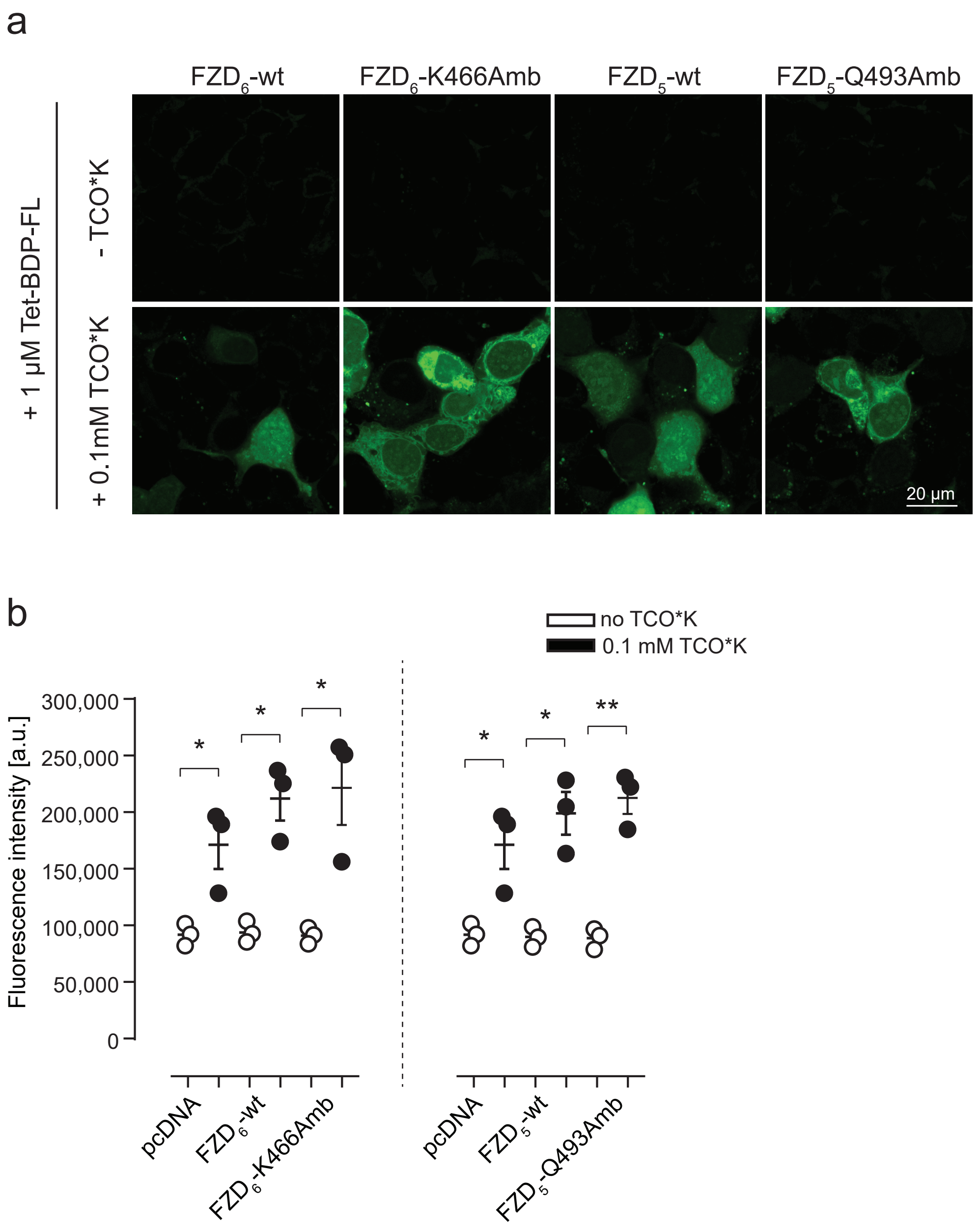

Supplementary Figure 4. **Fluorescence labeling of $FZD_5$ and $FZD_6$-TCO*K-incorporated mutants with Tet-BDP-FL.** a, Representative confocal images of HEK293T cells transiently cotransfected with $FZD_5$-wt, $FZD_6$-wt or the indicated amber mutants and the orthogonal tRNA/synthetase pair in absence (-) or presence (+) of 0.1 mM TCO*K. TCO*K-incorporated mutants were labeled with 1 μM of the cell membrane-permeable fluorescent dye Tet-BDP-FL. *Scale bar*, 20 μm. b, Fluorescence intensities of HEK293T cells transiently cotransfected with $FZD_5$-wt, $FZD_6$-wt or the indicated amber mutants and the orthogonal tRNA/synthetase pair in absence (-) or presence (+) of 0.1 mM TCO*K after labeling with Tet-BDP-FL measured in a plate reader assay. Data show mean ± s.e.m. of three individual experiments performed in triplicates. Results were analyzed with one-way ANOVA and uncorrected Fisher's LSD post-hoc test. Significance levels are given as * ($p < 0.05$), and ** ($p < 0.01$). *Amb*, amber mutant, *TCO*K*, TCO-Lysine, *Tet*, Tetrazine, *wt*, wild type.

### Supplementary Figure 5

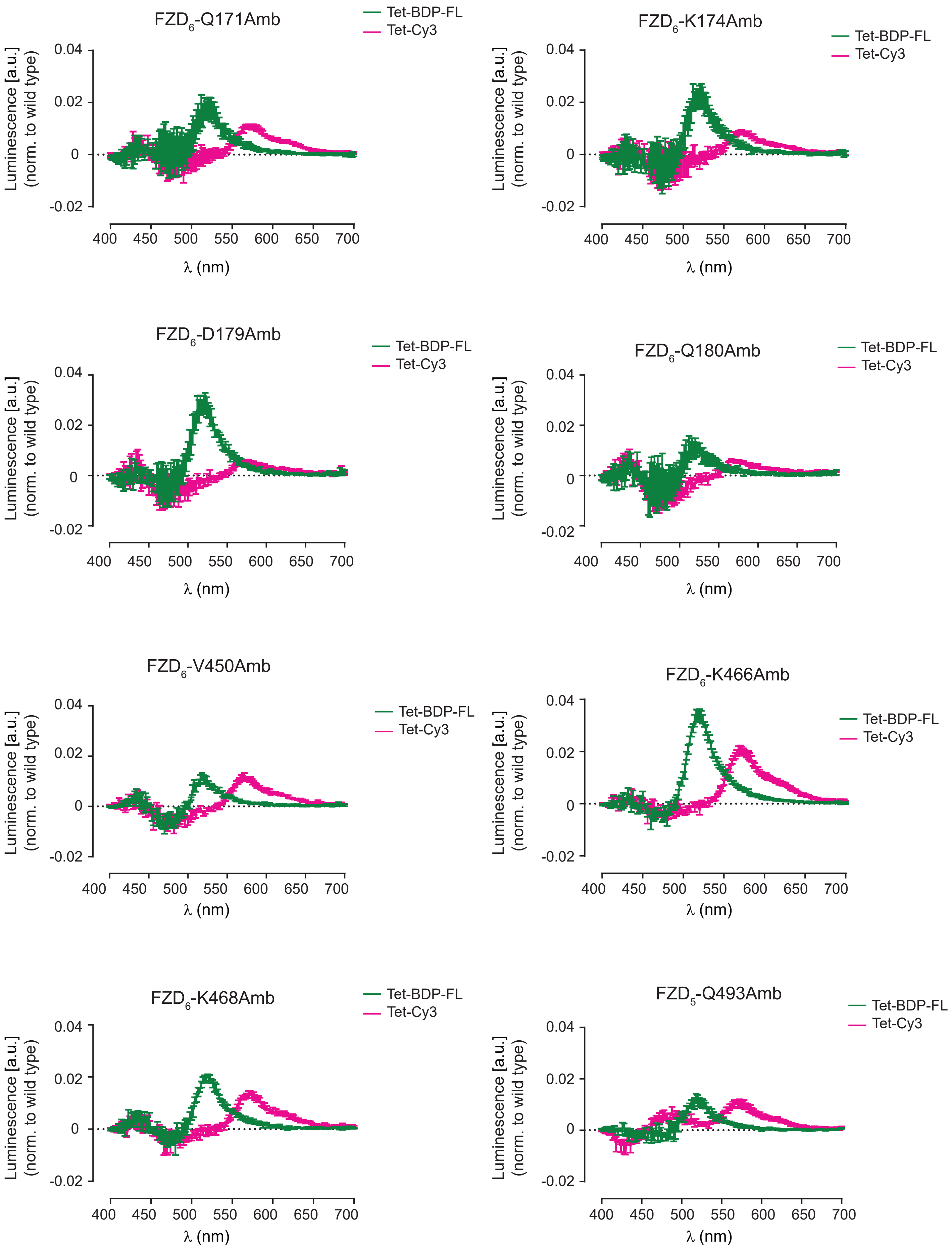

Supplementary Figure 5. **Comparison of different fluorescent dyes linked to $FZD_5$ and $FZD_6$ functioning as extracellular conformational BRET sensors.** Bioluminescence emission spectra of Nluc-tagged $FZD_5$ and $FZD_6$ TCO*K-incorporated mutant CRD sensors labeled with Tet-BDP-FL (green) or Tet-Cy3 (magenta) measured after addition of furimazine were normalized as a ratio of the maximal emission of each fluorescent protein. Bioluminescence obtained in $FZD_5$-wt or $FZD_6$-wt sensors (no TCO*K incorporated) was subtracted from the emission spectrum for each receptor mutant to correct for unspecific binding of the fluorescent dyes in three individual experiments. All experiments were performed in HEK293T cells cotransfected with the indicated $FZD_5$ or $FZD_6$ sensors and the orthogonal tRNA/synthetase pair. *Amb*, amber mutant, *Tet*, tetrazine.

### Supplementary Figure 6

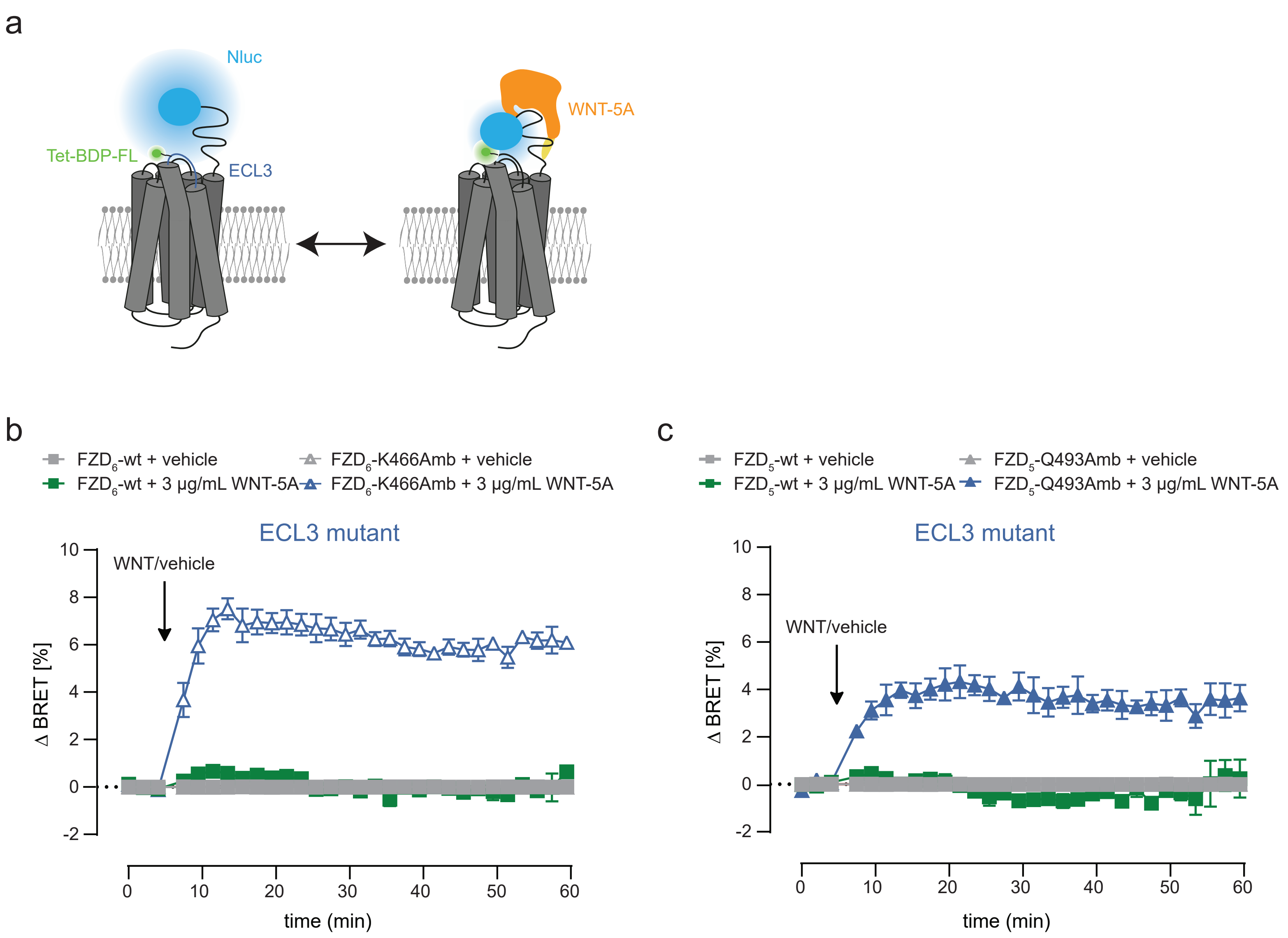

Supplementary Figure 6. **CRD movements in Tet-BDP-FL labeled $FZD_5$ and $FZD_6$ in response to WNT-5A.** a, Schematic depiction of the CRD sensor design with the N-terminally Nluc-tagged FZD. Fluorescence labeling of residues in the ECL3 (blue) region of the receptor with Tet-BDP-FL. b-c, BRET responses of $FZD_6$ (b) or $FZD_5$ (c) ECL3 amber mutant CRD sensors upon 3 µg/mL WNT-5A treatment or vehicle control. The arrow indicates the time point of WNT/vehicle addition. All experiments were performed in HEK293T cells cotransfected with the indicated $FZD_5$ or $FZD_6$ sensors and the orthogonal tRNA/synthetase pair. *Amb*, amber mutant, *ECL3*, extracellular loop 3, *Nluc*, NanoLuciferase, *Tet*, Tetrazine, *TCO*K*, TCO-Lysine, *wt*, wild type.

### Supplementary Figure 7

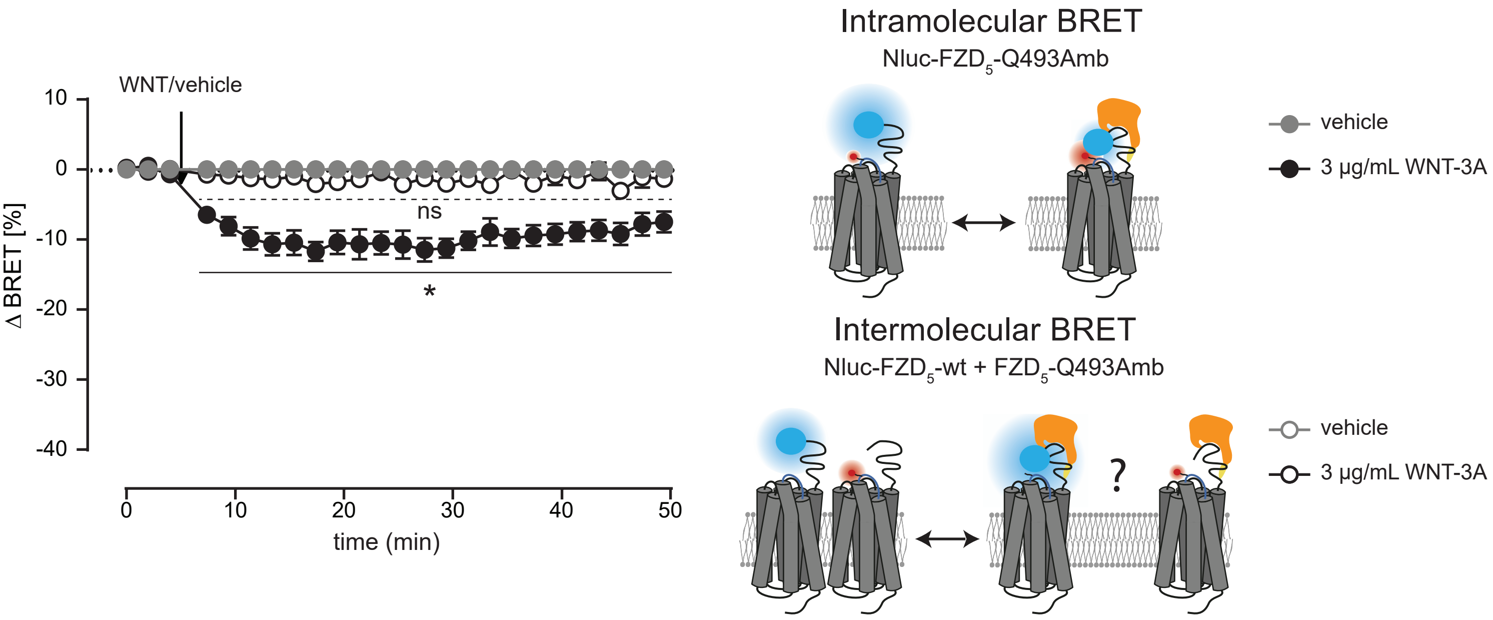

Supplementary Figure 7. **Intra- *versus* intermolecular BRET responses in $FZD_5$.** BRET responses of HEK293T cells expressing Nluc-$FZD_5$-Q493Amb (intramolecular BRET) in comparison to an intermolecular BRET control where Nluc-$FZD_5$-wt is cotransfected with a Nluc-lacking $FZD_5$-Q493Amb mutant upon 3 µg/mL WNT-5A treatment or vehicle control of four individual experiments. The arrow indicates the time point of WNT/vehicle addition. Differences between vehicle control and WNT-5A-induced BRET responses were analyzed with multiple t-test followed by Holm-Sidak multiple comparison. Significance levels are given as * ($p < 0.05$), and *ns* (not significant). *Amb*, amber mutant, *Nluc*, NanoLuciferase, *wt*, wild type.

### Supplementary Figure 8

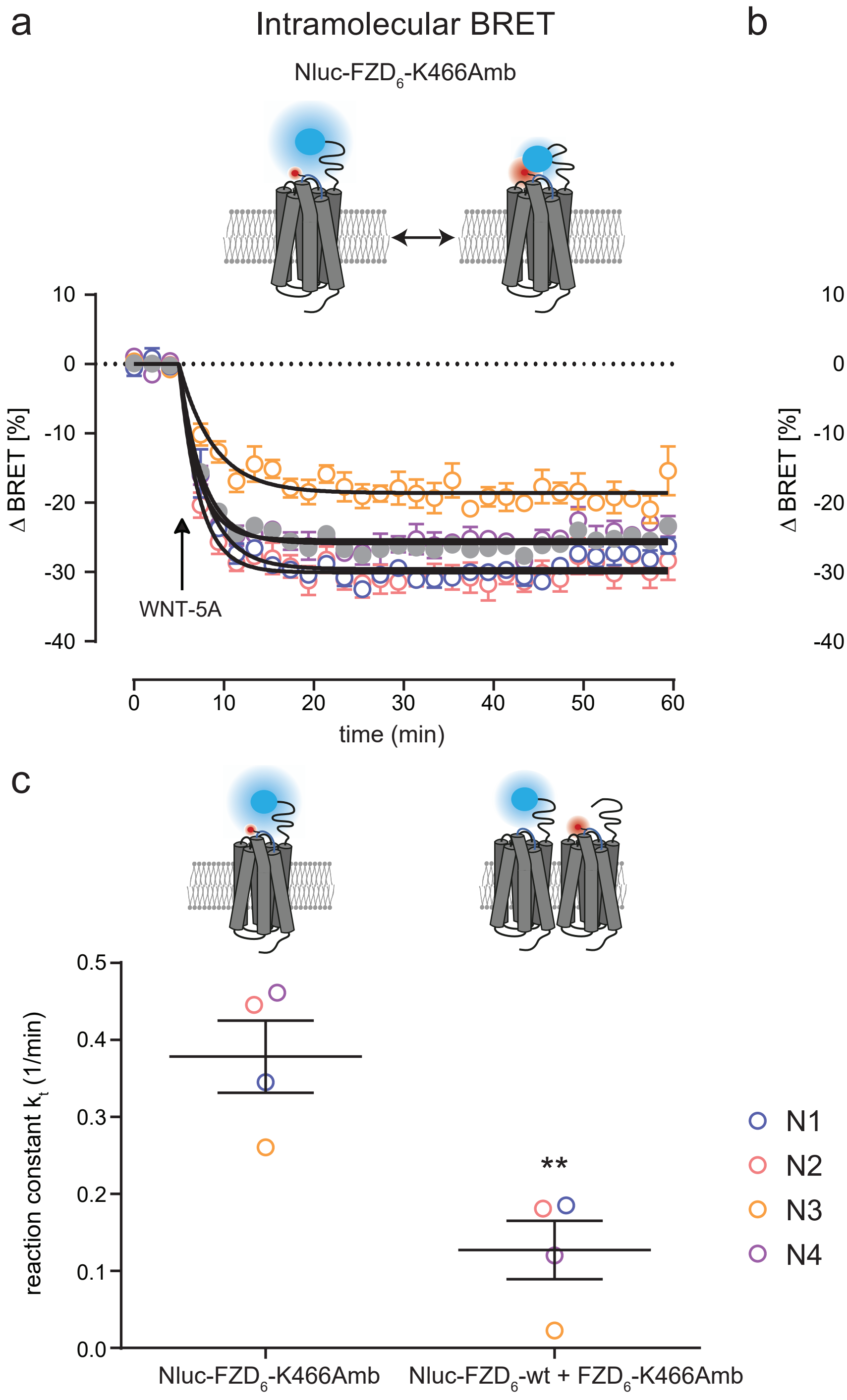

Supplementary Figure 8. **Intra- *versus* intermolecular BRET responses in $FZD_6$.** a, BRET responses of HEK293T cells expressing Nluc-$FZD_6$-K466Amb (intramolecular BRET) upon 3 µg/mL WNT-5A stimulation in four individual experiments (N1 – N4). b, BRET responses of HEK293T cells expressing an intermolecular BRET control where Nluc-$FZD_6$-wt was cotransfected with a Nluc-lacking $FZD_6$-K466Amb mutant upon 3 µg/mL WNT-5A treatment or vehicle control. c, Reaction constant k of $FZD_6$-K466Amb (intramolecular BRET) and $FZD_6$-wt + Nluc-lacking $FZD_6$-K466Amb intermolecular BRET control determined from fitted data in (a) and (b) by using the plateau followed by one phase decay equation. Differences of reaction constant k of the Nluc-$FZD_6$-K466Amb sensor and the Nluc-$FZD_6$-wt + $FZD_6$-K466Amb intermolecular BRET control were analyzed with Student's unpaired t-test. Significance levels are given as ** ($p < 0.01$). *Amb*, amber mutant, *Nluc*, NanoLuciferase, *wt*, wild type.

### Supplementary Figure 9

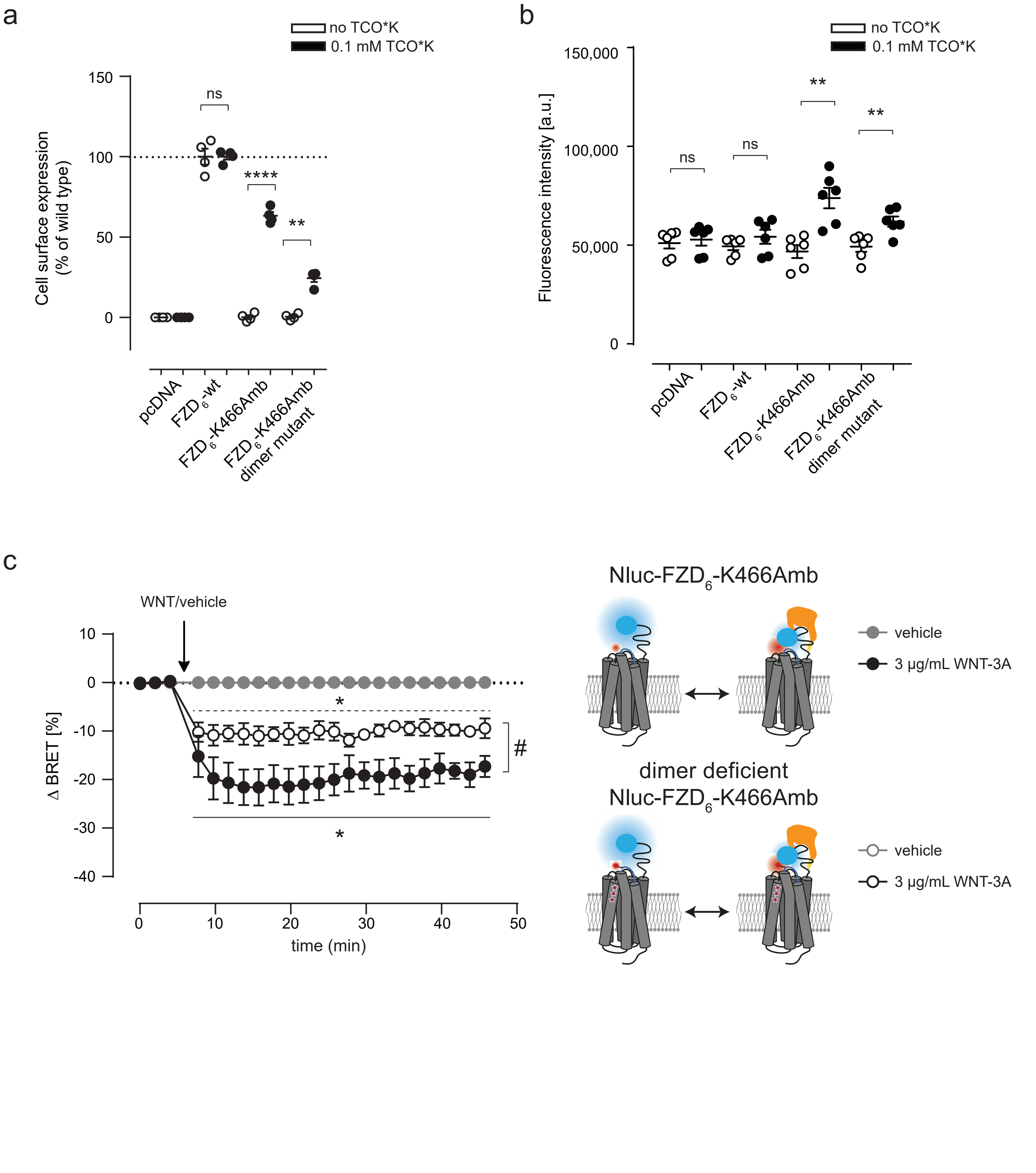

Supplementary Figure 9. **$FZD_6$-K466Amb dimer mutant controls for monomeric WNT-induced BRET responses.** a, Receptor surface expression of HEK293T cells transiently cotransfected with pcDNA3.1 or the indicated $FZD_6$ constructs with special attention on the $FZD_6$ dimerization deficient triple Ala mutant $FZD_6$-K466Amb-D365A/R368A/Y369A (named as $FZD_6$-K466Amb dimer mutant) and the orthogonal tRNA/synthetase pair in absence (-) or presence (+) of 0.1 mM TCO*K was quantified by whole-cell ELISA using an antibody against the N-terminal Nluc tag. Data show mean ± s.e.m. of four individual experiments performed in triplicates. Background fluorescence detected in pcDNA-transfected HEK293T cells was subtracted from all data, and mean values were normalized to wt $FZD_6$ surface expression. b, Fluorescence intensities of HEK293T cells transiently cotransfected with pcDNA3.1 or the indicated $FZD_6$ and the orthogonal tRNA/synthetase pair in absence (-) or presence (+) of 0.1 mM TCO*K after labeling with Tet-Cy3 measured in a plate reader assay. Data show mean ± s.e.m. of six individual experiments performed in triplicates. All results in (a) and (b) were analyzed with one-way ANOVA and uncorrected Fisher's LSD post-hoc test. Significance levels are given as ** ($p < 0.01$), **** ($p < 0.0001$), and *ns* (not significant). c, BRET responses of HEK293T cells expressing the Nluc-$FZD_6$-K466Amb dimer mutant upon 3 µg/mL WNT-5A stimulation in four individual experiments (N1 – N4). Differences between vehicle control and WNT-5A-induced BRET responses were analyzed with multiple t-test followed by Holm-Sidak multiple comparison. Significance levels are given as * ($p < 0.05$). Plateaus of WNT-5A induced BRET responses of the two different sensors were analyzed with extra sum of squares F test. Significance levels are given as # ($p < 0.05$). *Amb*, amber mutant, *Nluc*, NanoLuciferase, *TCO*K*, TCO-Lysine, *wt*, wild type.
